# Alphavirus infection triggers selective cytoplasmic translocation of nuclear RBPs with moonlighting antiviral roles

**DOI:** 10.1101/2021.10.06.463336

**Authors:** Wael Kamel, Vincenzo Ruscica, Azman Embarc-Buh, Zaydah R. de Laurent, Manuel Garcia-Moreno, Yana Demyanenko, Meghana Madhusudhan, Louisa Iselin, Aino Järvelin, Maximilian Hannan, Eduardo Kitano, Samantha Moore, Andres Merits, Ilan Davis, Shabaz Mohammed, Alfredo Castello

## Abstract

RNA is a central molecule for RNA viruses, acting as mRNA and genome. However, the interactions that viral (v)RNA establishes with the host cell is only starting to be elucidated. Here, we determine with unprecedented depth the composition of the ribonucleoproteins (RNPs) of the prototypical arthropod-borne Sindbis virus (SINV) using viral RNA interactome capture. We show that SINV RNAs engage with hundreds of cellular proteins and pathways, including a group of nuclear RNA-binding proteins (RBPs) with unknown roles in infection. Combining subcellular fractionation and proteomics with several orthogonal approaches, we demonstrate that these nuclear RBPs are selectively redistributed to the cytoplasm after infection, where they associate with the viral replication organelles. These nuclear RBPs potently supress viral gene expression, with activities spanning viral species and families. Our study provides a comprehensive and systematic analysis of SINV RNP composition, revealing a network of nuclear RBPs with moonlighting antiviral function.

**Research highlights:**

- SINV RNAs interact with over four hundred cellular RBPs
- SINV induces selective cytoplasmic translocation of a subset of nuclear RBPs
- These nuclear RBPs display potent antiviral effects
- The SF3B complex binds to SINV RNA and supresses infection in a splicing-independent manner

## INTRODUCTION

The alphavirus genus contains several arthropod-borne viruses (arboviruses) that can cause disease in humans, including chikungunya virus (CHIKV), Venezuelan equine encephalitis virus (VEEV), Ross River virus (RRV), Semliki Forest virus (SFV) and Sindbis virus (SINV) (Kurkela et al., 2004; Laine et al., 2004; Marks and Marks, 2016; Silva and Dermody, 2017). These arboviruses grow in the salivary glands of mosquitoes from the *Aedes* family and are transmitted to the human host through their bites (Lim et al., 2018). The habitats of the main vectors for alphaviruses, *Aedes aegypti and Aedes albopictus*, are expanding due global warming and international trade, raising concerns about the emergence of arboviral outbreaks and epidemics (Bonizzoni et al., 2013). Therefore, it is crucial to deepen our understanding of the host-virus interactions underpinning the infection of arthropod-borne viruses to identify potential targets for therapeutic intervention.

Alphaviruses have single stranded (ss), positive sense RNA genomes of ∼11kb that are capped and polyadenylated, resembling cellular mRNAs (Strauss and Strauss, 1994). Because of their limited coding capacity, they heavily rely on cellular proteins (Carrasco et al., 2018). Several studies have recently highlighted the importance of viral (v)RNA as a hub for crucial host-virus interactions (Iselin et al., 2022). vRNA assembles with viral and cellular RNA-binding proteins (RBPs) forming viral ribonucleoproteins (vRNPs) that are critical mediators and regulators of the viral lifecycle (Carrasco et al., 2018; Garcia-Moreno et al., 2019; Kim et al., 2020). Two proteome-wide approaches, viral crosslinking and solid-phase purification (VIR-CLASP) and crosslink-assisted messenger RNP purification (CLAMP) have been recently used to elucidate the composition of alphavirus RNPs (Iselin et al., 2021). VIR-CLASP focuses on the study of the interactions that CHIKV genomic (g)RNA establishes immediately after cell entry, revealing hundreds of early host-virus interactions (Kim et al., 2020). CLAMP focuses on the vRNAs synthesised after the burst of viral replication happening at later times post infection, and revealed many putative cellular interactors (Gebhart et al., 2020; Iselin et al., 2022; LaPointe et al., 2018). While informative, the CLAMP datasets showed low incidence of *bona fide* RBPs due to technical limitations that are discussed in detail elsewhere (Iselin et al., 2021). New, more robust approaches are therefore required to profile the post-replicative alphaviral RNPs.

By applying a proteome-wide approach called viral RNA interactome capture (vRIC) (Kamel et al., 2021), we discovered that SINV RNAs interact with over four hundred cellular RBPs. SINV RNPs exhibit notable defences over cellular RNPs, including a striking enrichment in kinases that can control the replication microenvironment and a differential set of translation initiation factors. We also report a group of nuclear RBPs that is translocated from the nucleus to the cytoplasm upon infection, where they associate with SINV RNA and/or its RNA-dependent RNA polymerase (RdRp) complex. Functional assays revealed that these nuclear RBPs potently supress viral replication, suggesting that they are part of an uncharacterised antiviral protein network. We discovered that the SF3B complex, which is part of the U2 small nuclear ribonucleoprotein (snRNP) of the spliceosome, associates with SINV RNA and represses viral gene expression in a splicing-independent manner.

## RESULTS

### Elucidating the composition of SINV RNPs

SINV infection causes a profound remodelling of the cellular RNA-binding proteome (RBPome) (Garcia-Moreno et al., 2019). This implies that the pool of cellular RBPs that SINV RNAs encounter varies as infection progresses. To comprehensively profile the complement of cellular RBPs that interact with SINV RNA in SINV-infected cells after the replication burst, we used viral RNA interactome capture (vRIC) (Kamel et al., 2021). Briefly, cells were infected with SINV and then treated with 4-thiouridine (4SU) and an inhibitor of cellular RNA polymerases (Figure 1A). Because the viral RdRps are refractory to cellular RNA polymerase inhibitors, 4SU is primarily incorporated into newly synthesised vRNA. Zero distance crosslinking between RBPs and vRNA are achieved by activation of the 4SU molecules with ultraviolet (UV) light at 365 nm. Natural bases have negligible absorption at this wavelength limiting protein-RNA crosslinking to 4SU-containing RNAs (Baltz et al., 2012; Castello et al., 2012). Upon lysis under denaturing conditions, proteins covalently bound to vRNA are isolated with oligo(dT) magnetic beads under stringent denaturing washes, eluted by heat and RNase treatment, and analysed by quantitative proteomics.

**Figure 1.**
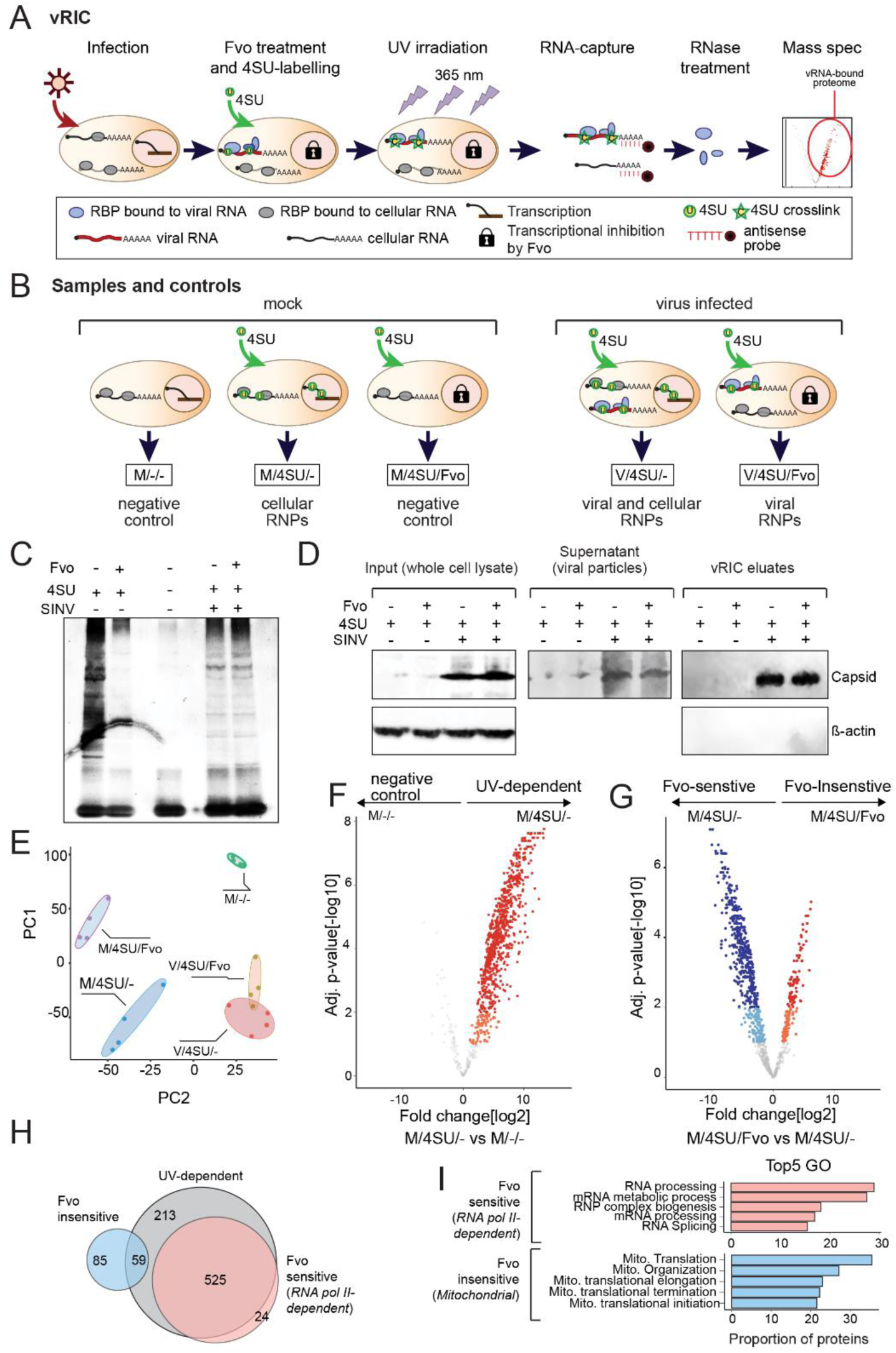
vRIC implementation in SINV infected cells. A) Schematic of the viral RNA interactome capture (vRIC) workflow. B) Schematic of the different experimental conditions and controls used in the study. C) Sliver staining analysis of vRIC eluates. C) Immunoblotting of SINV capsid in whole cell lysates, supernatant and vRIC eluates, using β-actin as control. E) Principal component analysis (PCA) of vRIC experimental and controls samples after proteomic analysis F and G) Volcano plots showing the fold change (log_2_) and adjusted p-value of each protein (dot) in the M/-/- versus M/4SU/- (F) and M/4SU/- versus M/4SU/Fvo (G). Proteins enriched within 1% FDR are coloured in red and blue and those within 10% FDR in orange and cyan. H) Venn diagram showing the overlapping between the significantly enriched proteins in the comparisons shown in F-G. These groups are classified: UV-dependent RBPome (M/-/- vs M/4SU/-), Fvo-sensitive (enriched in M/4SU/- over M/4SU/Fvo, cRNPs) and Fvo-insensitive (enriched in M/4SU/Fvo over M/4SU/-, mitoRNPs). I) Top enriched Gene Ontology (GO) terms for Molecular processes in Fvo-sensitive and Fvo-insensitive RBP sets. Related to Figure S1.

Initially, we tested the use of the actinomycin D (ActD), 5,6-dichloro-1-beta-D-ribofuranosylbenzimidazole (DRB) and flavopiridol (Fvo) to inhibit cellular RNA polymerases (Figure S1A). (Garcia-Moreno et al., 2019), we observed that neither DRB nor Fvo had detrimental effects on SINV_mCherry_ fitness (Figure S1A). Conversely, ActD caused a substantial reduction in viral gene expression. Inhibition of the cellular RNA polymerase II was tested in a cell line expressing eGFP from a strong cytomegalovirus promoter inducible by deoxycycline (Dox). ActD and, particularly, Fvo strongly suppressed eGFP expression, whereas DRB had a mild effect (Figure S1B). We selected Fvo for further experiments given that it strongly inhibits RNA polymerase II dependent transcription while not having effects in SINV fitness. Next, we optimised the 4SU pulse to maximise incorporation into vRNA without affecting replication. When SINV_mCherry_ and 4SU were added simultaneously, viral gene expression was inhibited in a dose-dependent manner (Figure S1C and D). The negative effects of 4SU were prevented by delaying its addition for 2h (Figure S1E). Lack of 4SU-dependent inhibitory effects in viral fitness when added at 2 hours post infection (hpi) was also observed for SFV (Figure S1F) and SARS-CoV-2 (Kamel et al., 2021).

To elucidate the composition of SINV RNPs, we applied the optimised vRIC protocol to SINV-infected cells, followed by quality control analysis and proteomics (Figure S2A-E). We included all the controls specified in Figure 1B. In the absence of 4SU and virus (M/-/-), the overall protein intensity observed was very low, leading to a remarkably “clean” lane in a silver staining (Figure 1C, third lane and S2F). Conversely, addition of 4SU and UV_365_ (M/4SU/-) led to the isolation of a complex protein pool that resembled previously established RBPomes (Figure 1C, first lane and S2F) (Baltz et al., 2012; Castello et al., 2012). Importantly, addition of Fvo (M/4SU/Fvo) strongly reduced protein isolation, reflecting a lack of 4SU incorporation into poly(A) RNAs (Figure 1C, second lane and S2F). Indeed, principal component analysis (PCA) revealed that M/4SU/- and M/4SU/Fvo samples cluster separately (Figure 1E).

To generate a reference group for the vRNPs, we defined the composition of cellular RBPs that bind cellular poly(A)+ RNA by analysing the uninfected samples with and without 4SU (M/4SU/- vs M/-/-). The proteomic analysis revealed 797 proteins significantly enriched in a UV-dependent manner, with most of the identified proteins being *bona fide* RBPs (Figure 1F, S2G and Table S1). We noticed that a small set of 144 RBPs was enriched in Fvo-treated uninfected samples over its untreated control (M/4SU/Fvo vs M/4SU/-), which was primarily composed by ‘mitochondrial’ proteins (Figure 1G-I, Table S3). These proteins probably derive from the isolation of mitochondrially encoded poly(A)+ RNAs by the lack of inhibition of mitochondrial RNAP by Fvo (Barchiesi and Vascotto, 2019; Dhir et al., 2018; Temperley et al., 2010). For simplicity, we refer to these Fvo-insensitive proteins as “mitochondrial (mito)RNPs”, and the Fvo-sensitive (RNA polymerase II-dependent) proteins as “cellular (c)RNPs” (Figure S2I, Table S2 and S3).

In contrast to uninfected cells, a rich protein pattern was observed in SINV-infected cells despite the presence of Fvo (Figure 1C, lane 5). SINV capsid was highly enriched, as expected from a viral RBP (Figure 1D) (Garcia-Moreno et al., 2019; Sokoloski et al., 2017). Fvo treatment induced a reduction in protein content in eluates from SINV infected cells (V/4SU/Fvo vs V/4SU/-, Figure S2H), which is consistent with a Fvo-dependent depletion of RBPs interacting with cellular mRNAs when the RNA polymerase II is inhibited.

### Similarities and divergences between cellular and SINV RNPs

vRIC identified 459 RBPs (345 with 1% FDR and 114 with 10% FDR) enriched in SINV-infected samples (V/4SU/Fvo) over the mock control (M/4SU/Fvo) (Figure 2A, Table S4). Interestingly, cRNPs and vRNPs shared over 50% of their components, while nearly no overlap was observed with mitoRNPs (Figure 2B and S2I). These results indicate that mitoRNPs are sub-stoichiometric and only observable when cRNPs and vRNPs are absent. We found that nearly 50% of the proteins in the vRNP and cRNP datasets harbour well-established RNA-binding domains (RBDs) (Figure 2C), while mitoRNP components and proteins enriched only in vRNPs (vRNPen) have a higher incidence of RBPs with RBDs that are non-canonical or uncharacterised (Figure 2C and S2J). Intrinsically disordered regions (IDRs) are common across RBPs and have prevalent roles in RNA binding (Castello et al., 2012; Castello et al., 2016; Jarvelin et al., 2016). The proportion of IDRs within vRNP and RNP components was similar, unlike mitoRNPs, which are known to be depleted in IDRs (Figure 2D and S2K) (Fukuchi et al., 2011). Altogether, these results indicate that RBPs within vRNP and cRNP share important biochemical properties and differ from mitochondrial RBPs. To understand the differences of the RBPs within vRNPs and cRNPs, we analysed their binding profiles when available using the ENCODE eCLIP database (Van Nostrand et al., 2016). vRNPs and cRNPs include 140 RBPs with available eCLIP data (i.e., their footprints on cellular RNAs). We grouped them based on their binding preferences for different regions of cellular mRNAs and found that vRNPs are enriched in RBPs that typically bind to 5’ and 3’ untranslated regions (UTRs), while cRNPs have a higher proportion of RBPs that bind to introns (Figure 2E). This compositional bias highlights the importance of the 5’ and 3’ UTRs of SINV RNAs in replication, viral gene expression and RNA packaging (Carrasco et al., 2018; Strauss and Strauss, 1994). The lack of introns in SINV RNAs explains the relative depletion of intron-binding proteins.

**Figure 2.**
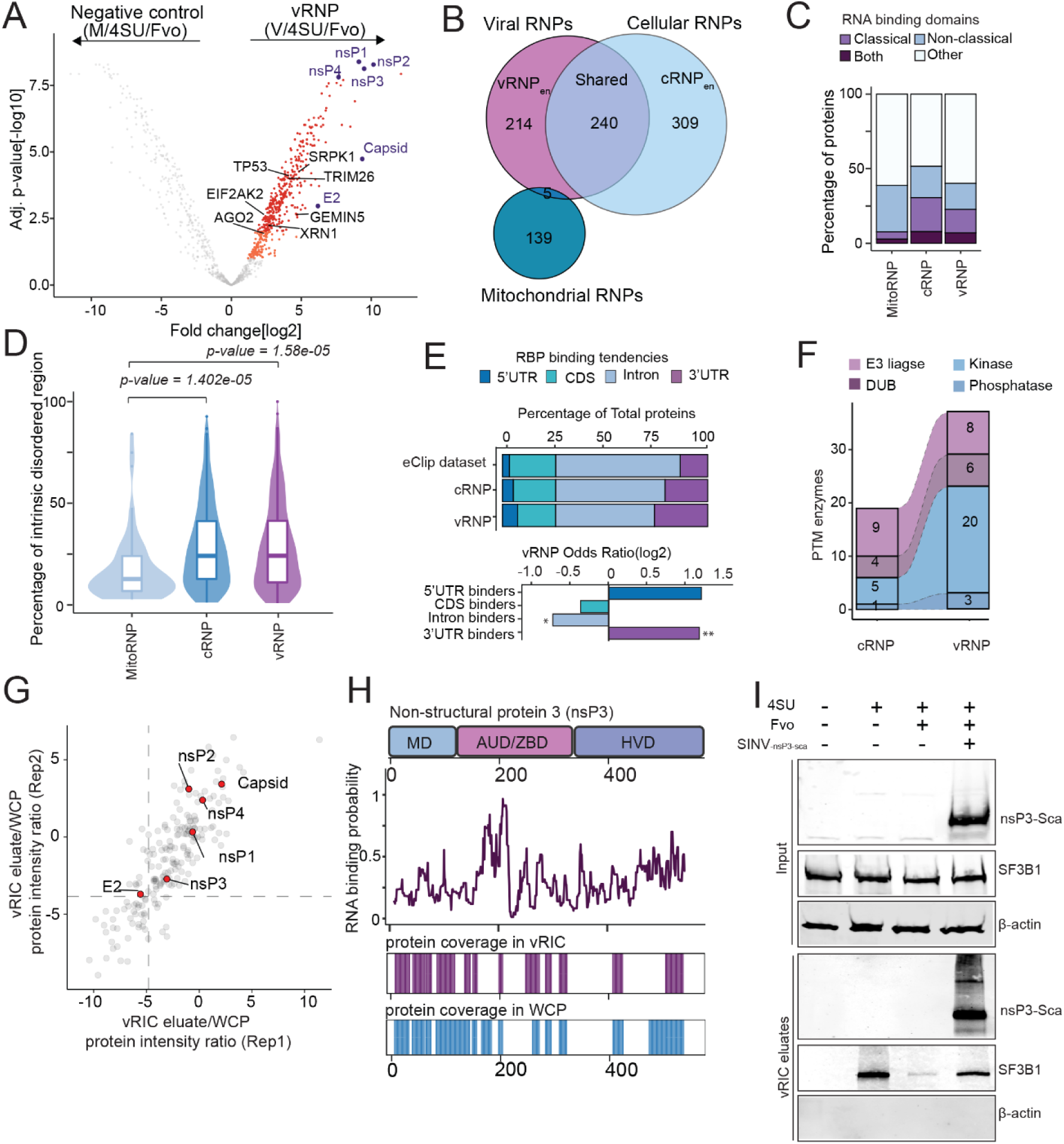
Uncovering the composition of SINV RNPs. A) Volcano plot showing fold change (log_2_) and adjusted p-value for each protein (dot) in the comparison between M/4SU/Fvo versus V/4SU/Fvo. Proteins enriched within 1% FDR are coloured in red and those within 10% FDR in orange. Proteins significantly enriched in the V/4SU/Fvo are classified as SINV RNP components. B) Venn diagram comparing the components of vRNPs, cRNPs and mitoRNPs. C) Bar plot defining the proportion of RBPs with classical, non-classical, both or undefined RBDs in components of vRNPs, cRNPs and mitoRNPs. D) Prevalence of intrinsic disorder regions (IDRs) in vRNPs, cRNPs and mitoRNPs. p-values were estimated by Welch’s t-test. E) Upper panel, bar plot showing the proportion of RBPs with biases towards 5’ UTR, 3’ UTR, CDS and introns in vRNPs, cRNPs and the whole ENCODE eCLIP superset. Lower panel, incidence of RBPs with preferences for intron, 5’ and 3’UTR binders in vRNPs over cRNPs. *, p <0.1 and **, p< 0.05 estimated by Fisher’s Exact test. F) Bar plot defining the number of RBPs classified as post-translational modifying enzymes (only showing E3ligases, deubiquitinases, kinases and phosphatases) in cRNPs and vRNPs. G) Scatter plot showing the protein intensity ratios in vRIC eluate over whole cell proteome (WCP) in two biological replicates. Viral proteins detected by vRIC are highlighted. H) Upper panel, domain architecture of nsP3 (MD, macro domain; AUD/ZBD, alphavirus-unique or zinc-binding domain, and HVD, hypervariable domain). Middle panel, profile showing the probability for RNA binding at every amino acid of nsP3. Lower panels, position of the peptides identified in vRIC eluates and WCP. I) Immunoblotting analysis of vRIC inputs (whole cell lysates), and eluates of SINV_nsP3-mScarlet_ infected cells. nsP3-mScartlet, nsP3-Sca Related to Figure S2.

Another interesting feature of vRNPs is the enrichment of post-translational modification (PTM) enzymes over cRNPs (Figure 2F). This phenomenon was predominantly driven by kinases, including important regulators of infection such as the antiviral EIF2AK2 (PKR) and PRKDC (Hull and Bevilacqua, 2016; Patricio et al., 2022; Peters et al., 2013), and the ‘pro-viral’ SRPK1 (Fukuhara et al., 2006; Garcia-Moreno et al., 2019). Beyond these expected kinases, we noticed the presence of several mitogen-activated protein kinases (MAPKs), and cyclin-dependent kinases.

Interestingly, we also noticed a differential set of eukaryotic translation initiation factors interacting with cellular and SINV RNAs. SINV subgenomic (sg)RNA is capped and polyadenylated but its translation follows a non-canonical cap-dependent mechanisms with unknown mechanism (Carrasco et al., 2018; Castello et al., 2006; Garcia-Moreno et al., 2013; Ventoso et al., 2006). Interestingly, the cap-binding protein EIF4E, the helicase EIF4A2, the Met-tRNAi delivery factor EIF2S1 (EIF2α) and EIF2S2 (EIF2β) were not detected in the vRNP, while being present in cRNPs (Figure S2M). By contrast, EIF4G1, EIF4G2, EIF4A1, EIF4B and several eIF3 components were detected in both vRNPs and cRNPs. Altogether, our results reveal differences between SINV RNPs and cRNPs that deserve further investigation.

### Unveiling the SINV proteins that interact with vRNA

vRIC identified six viral proteins within vRNPs, which include the non-structural proteins (nsP)1, nsP2, nsP3 and nsP4 and the structural proteins capsid and E2 (Figure 2A, S2L and Table S4). To determine whether the capture of these proteins truly reflect a *bona fide* interaction with vRNA and their identification is not a product their high abundance, we normalised their intensities in vRIC eluates to those in the whole cell proteome, generating a ‘UV_365_ crosslink-ability’ score (Figure 2G). Capsid displayed the highest crosslink-ability, as expected from its involvement in viral RNA packaging (Sokoloski et al., 2017), followed by the capping enzyme nsP1, the helicase nsP2 and the RdRp nsP4. We found here that nsP3 also interacts with SINV RNAs (Figure 2G), although its RNA-binding activity remains poorly understood (Gotte et al., 2018). To validate nsP3 RNA-binding activity, we first used a RBDetect software, which predicts RNA-binding regions based on sequence similarities with RBDs in human RBPs (Hobor et al., 2018; Kamel et al., 2021). RBDetect revealed a high-confidence RNA-binding site placed at the AUD domain (Figure 2H), which is consistent with previous data on CHIKV nsP3 (Gao et al., 2019). Additionally, we noticed that the nsP3 peptide coverage was high in vRIC eluates, strongly supporting its interaction with vRNA (Figure 2H). To validate these results, we next infected cells with a chimeric virus expressing mScarlet fused to nsP3 (SINV_-nsP3-Scarlet_) and performed a vRIC experiment. Interestingly, nsP3-mScarlet was enriched in eluates of infected cells (Figure 2I), showing that nsP3 does not only interact with vRNA as part of the nsP1-4 polyprotein, but also as individual polypeptide. Nevertheless, nsP3 crosslinking-ability was substantially lower than that of the other nsPs (Figure 2G), suggesting a more transitory or spatially suboptimal interaction with RNA. Our data suggests that E2 also crosslinks to vRNA, in agreement with previously results (Garcia-Moreno et al., 2019). However, E2 crosslinking efficiency is the lowest of all SINV proteins indicating low affinity and very transitory interaction with vRNA (Figure 2A and G), in analogy with the spike of SARS-CoV-2 (Kamel et al., 2021).

### Uncovering the nsP3 interactome as a proxy for replication-associated vRNPs

Alphavirus replication takes place at bulb-like, membrane-bound vesicles known as the replication organelles (ROs), in which the RdRp complex (nsP1-4) and vRNA accumulates. Recent Cryo-EM structure of alphaviral ROs suggests that the location nsP3 is at the outer rim together with its interacting host proteins (Tan et al., 2022). Single molecule fluorescence *in situ* hybridisation (smFISH) and immunofluorescence using SINV_-nsP3-mScarlet_ showed that nsP3-mScarlet and SINV RNA accumulate together in cytosolic foci that are compatible with ROs (Figure 3A). To identify the protein partners of the RdRp, we used the nsP3-mScarlet produced by SINV_-nsP3-mScarlet_ as bait for immunoprecipitation (IP). nsP3-mScarlet was strongly enriched in the eluates, and led the co-precipitation of a discrete set of interactors that were absent in the negative control (Figure 3B-C). Proteomic analysis revealed 378 interactors of nsP3 across a wide range of fold changes and p-values. Amongst the most prominent interactors of nsP3, we identified the RdRp components nsP1, nsP2 and nsP4 and the cellular proteins G3BP1 and 2 (Kim et al., 2016) (Figure 3D, S3A-B and Table S5). Because viral proteins can potentially be identified due to their high abundance in infected cells, we normalised their intensities in the eluates to those in the whole cell lysate. This revealed a specific enrichment of nsP1, nsP2 and nsP4, confirming that the IP enriches for the RdRp complex (Figure 3E). Several ADP-ribosylation factors were detected in nsP3 IP, including the poly-ADP-ribose polymerase PARP1, and ARF3, ARF4, ARF5 and ARL6IP4. This aligns well with the known role of nsP3 at suppressing and reversing ADP ribosylation through its macrodomain (Gotte et al., 2018; Park and Griffin, 2009). Importantly, 25% of the nsP3 interactors also bind to SINV RNAs (i.e. vRNP dataset, Figure 3F), with these cellular RBPs being in average the most enriched proteins in the IP implying that they are the central partners of the RdRp complex (Figure 3G). Moreover, many of the nsP3 interactors have been functionally linked to vRNA metabolism and to the localization of proteins to membrane-bound organelles, which is compatible with processes occurring within the ROs (Figure 3H). The nsP3 interactome thus confirms that many of the proteins identified by vRIC in Figure 2A associate viral replication centres and, consequently, are proximal to vRNA, validating our results by an orthogonal approach. Moreover, it provides evidence for many proteins of functions related to replication or immediately downstream steps of viral gene expression.

**Figure 3.**
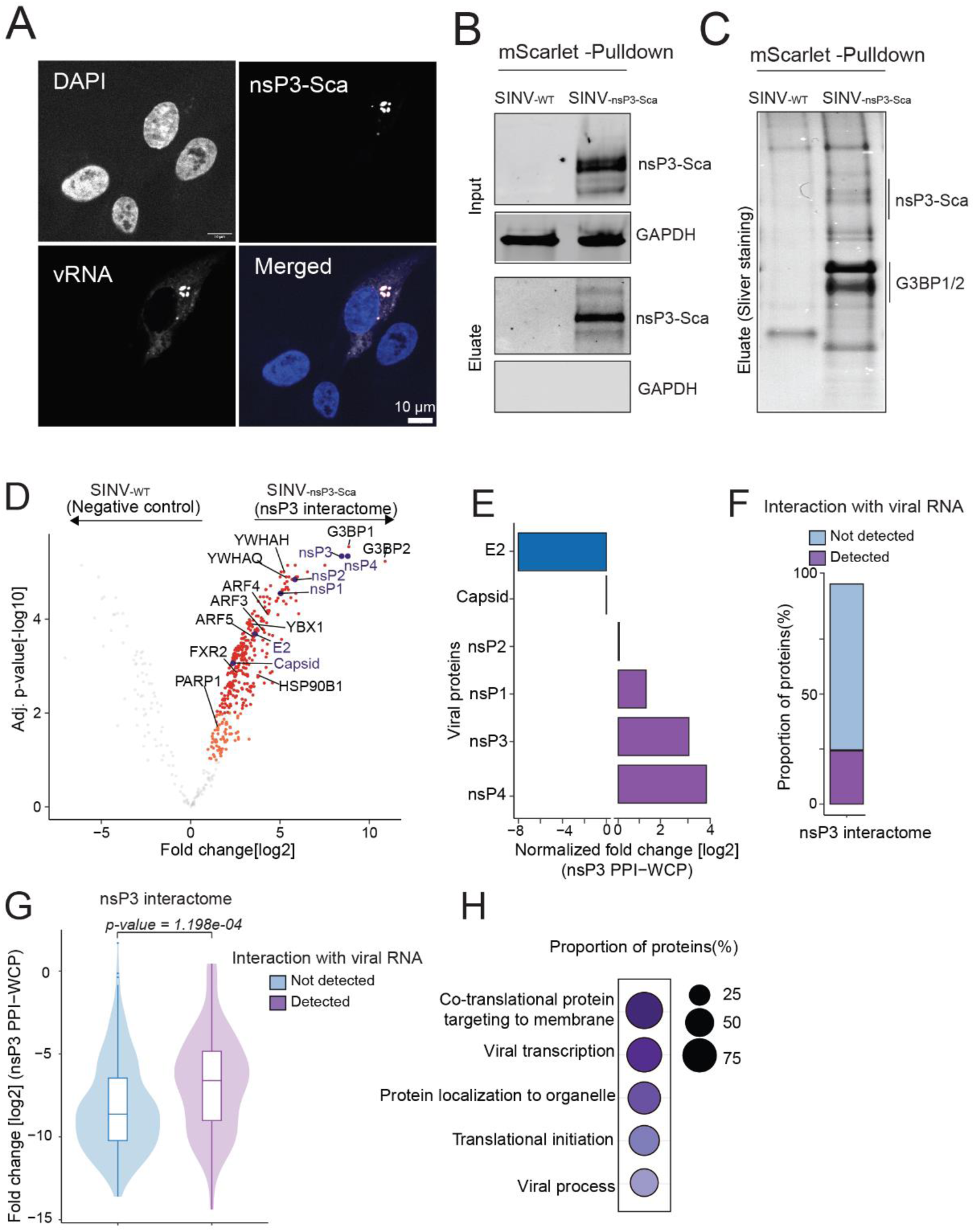
Viral protein nsP3 interactome in SINV-infected cells. A) smFISH and fluorescence analysis of SINV_nsP3-mScarlet_-infected A549 cells, showing viral RNA, nsP3-mScarlet (nsP3-Sca) and DAPI signal. B) Immunoblotting analysis of the immunoprecipitation with RFB nanobodies in extracts from SINV and SINV_nsP3-mScarlet_ infected cells. C) Sliver staining of the eluates of the nsP3-Sca immunoprecipitation. D) Volcano plot showing fold change (log_2_) and adjusted p-value for each protein (dot) in the immunoprecipitation in extracts from SINV and SINV_nsP3-mScarlet_ infected cells. Proteins enriched within 1% FDR are coloured in red and those within 10% FDR in orange. E) Normalized fold change (log2) of viral protein intensity ratios in the eluates of SINV nsp3-Sca co-immunoprecipitation versus whole cell lysate. F) Bar plot highlighting the proportion of viral RNA interactors that also binds to nsP3 G) Distribution of proteins intensities ratios in the eluates of nsp3-Sca immunoprecipitation versus whole cell lysate for nsP3 interactors that also bind (detect) or not (not detected) to SINV RNAs. p-values were estimated by Welch’s t-test. H) GO enrichment of cellular processes for RBPs that interact with nsP3 and SINV RNA over total SINV RNA interactome. Related to Figure S3.

### Specific cytosolic translocation of a set of nuclear RBPs that interact with SINV RNAs

Although SINV is a cytoplasmic virus, we noticed that a substantial proportion of the RBPs that bind to SINV RNAs are nuclear (Figure 4A). To test if this interaction takes place in the cytoplasm, we performed nuclear/cytoplasmic fractionation. After fraction quality analysis, samples were analysed by proteomics (Figure 4B and S4A). PCA confirmed that nuclear and cytoplasmic fractions differ in composition (Figure S4B). The nuclear/cytoplasmic ratios of most cellular proteins were unaltered by SINV infection (Figure 4C, left panels), which demonstrates that nuclear envelope integrity is maintained upon SINV infection. When focusing only on the components of SINV RNPs, we observed that nuclear RBPs exhibited a striking increase in cytoplasmic localisation (Figure 4C, right panels). These include the few RBPs known to relocate to the cytoplasm upon alphavirus infection (Figure S4C) (Lloyd, 2015; Sanz et al., 2015).

**Figure 4.**
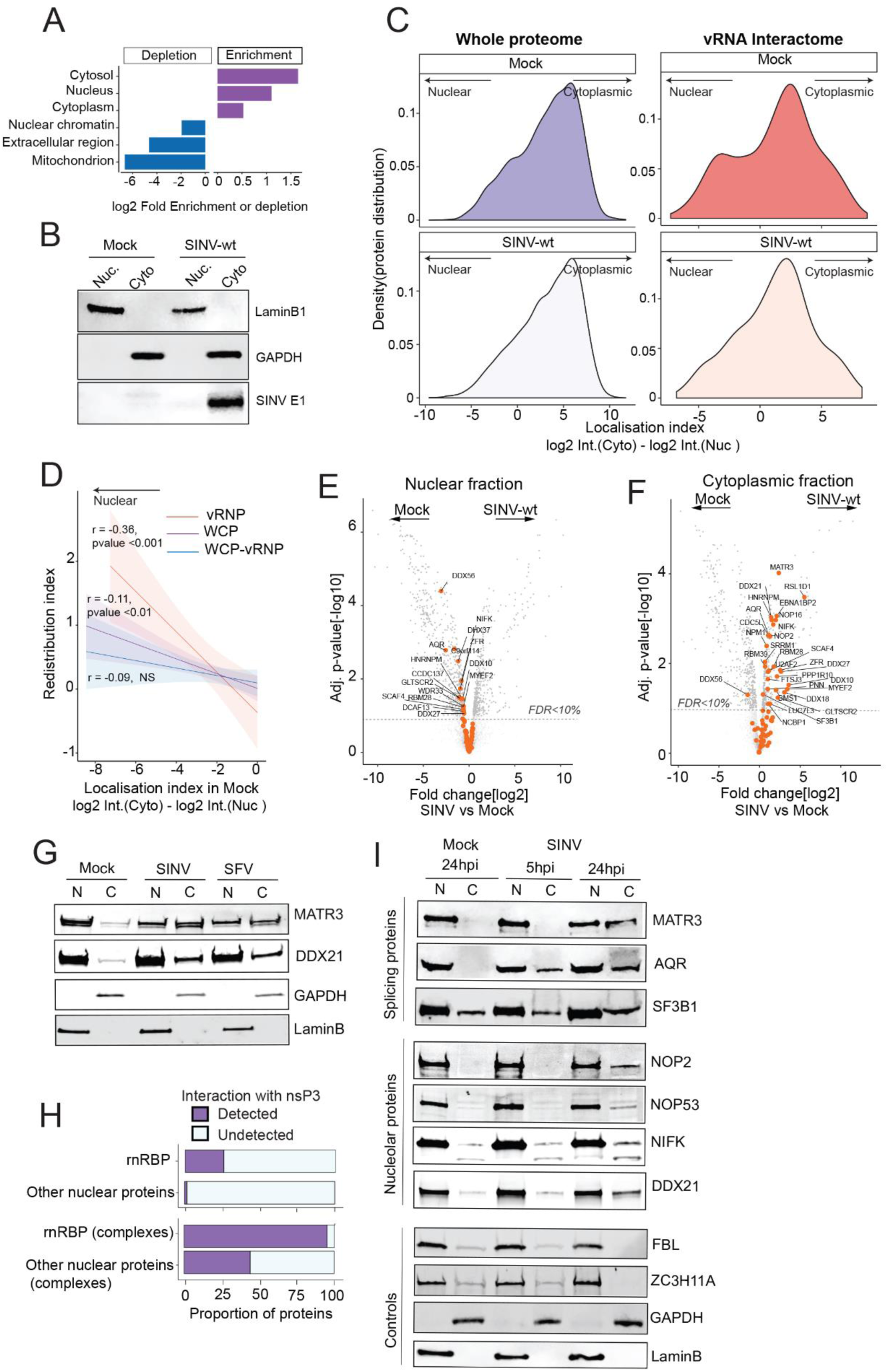
Nucleo-cytoplasmic redistribution of nuclear vRNP components. A) Bar plot showing ‘cellular component’ GO terms related to subcellular localisation that are enriched in vRNPs over the human proteome. B) Quality control Western blot analysis for the nucleo-cytoplasmic fractionation experiments in mock and SINV-infected cells. C) Kernel density analysis of the protein localisation index, which estimated as the [log2] ratio of protein intensity in the cytoplasmic fraction over the nuclear fraction. Density plots are shown for the proteins within the whole cell proteome (WCP) and vRNPs in both mock-infected and SINV-infected cells. D) Linear regression analysis of the localisation index against the redistribution index (i.e. localisation index of each protein in SINV infected vs mock cells) for the proteins within WCP, vRNPs, and WCP without vRNPs components (related to Figure S3D) E-F) Volcano plots showing the fold change and adjusted p-value for each protein in the nuclear (E) and cytoplasmic fractions (F) of SINV- and mock-infected cells. G) Bar plots defining the proportion of redistributed nuclear (rn)RBPs (upper panel) and their associated protein complexes (lower panel) in comparison to other nuclear proteins and their protein complexes detected in SINV nsP3 interactome. Curated human complexes were obtained from CORUM database (Giurgiu et al., 2019). H) Immunoblotting analysis of the nuclear and cytoplasmic fraction of SINV and SFV infected (24hpi) and uninfected cells. I) Immunoblotting of the representative proteins from (E-F), in the nuclear and cytoplasmic fraction of SINV- infected (5 and 24hpi) and uninfected cells. FBL (Fibrillarin), ZC3H11A, laminB1 and GAPDH were used as controls for nucleolar, nucleoplasmic, nuclear envelope and cytoplasmic proteins. Related to Figure S4.

To assess the magnitude and directionality of the changes in subcellular localisation, we generated a redistribution index for each protein (i.e. cytoplasmic/nuclear protein intensity ratio in mock versus infected cells) (Figure S4D). For simplicity, cellular proteins were grouped into nucleus-enriched or cytoplasm-enriched. Interestingly, most of the proteins in the cellular proteome with high redistribution index are nuclear in uninfected cells (r= −0.117, p <0.01), suggesting that the directionality of the translocation in infected cells is nucleus-to-cytoplasm and not the opposite (Figure 4D, S4E). This redistribution bias became more dominant when focusing exclusively in vRNP components (r= −0.35, p <0.001) (Figure 4D, S4E and F). Indeed, subtraction of the vRNP components from the whole proteome reduced the redistribution bias to non-significant (r=-0.056) (Figure 4D). No significant changes in the redistribution index were observed for the proteins enriched in cRNP (cRNPen; r=-0.09) (Figure S4E). These findings thus imply that vRNA is the main source of changes in subcellular localisation, probably recruiting and retaining cellular RBPs.

Nuclear vRNPs that show i) cytoplasmic-redistribution (Figure S4E, mid panel) and ii) differential enrichment in nucleus or/and cytoplasm after SINV infection (Figure 4E) were selected and assigned as ‘redistributed nuclear (rn)RBPs’. Interestingly, most of the rnRBPs are splicing factors or nucleolar proteins (Figure S4G, Table 1). Many of rnRBPs interact with the RNA of different cytoplasmic viruses (Figure S4H), suggesting that translocation of these rnRBPs to the cytoplasm may be common in cells infected with cytoplasmic RNA viruses. Indeed, we observed similar nucleo-cytoplasmic shuttling for MATR3 and DDX21 during SFV infection (Figure 4G). The nsP3 interactome is relatively depleted of nuclear proteins and their complexes [extracted from (Giurgiu et al., 2019)] (Figure S4I); however, it is enriched in rnRBPs and their protein partners (Figure 4H). These results highlight vRNA and proteins as potential ‘recruiters’ of rnRBPs.

**Table 1.**
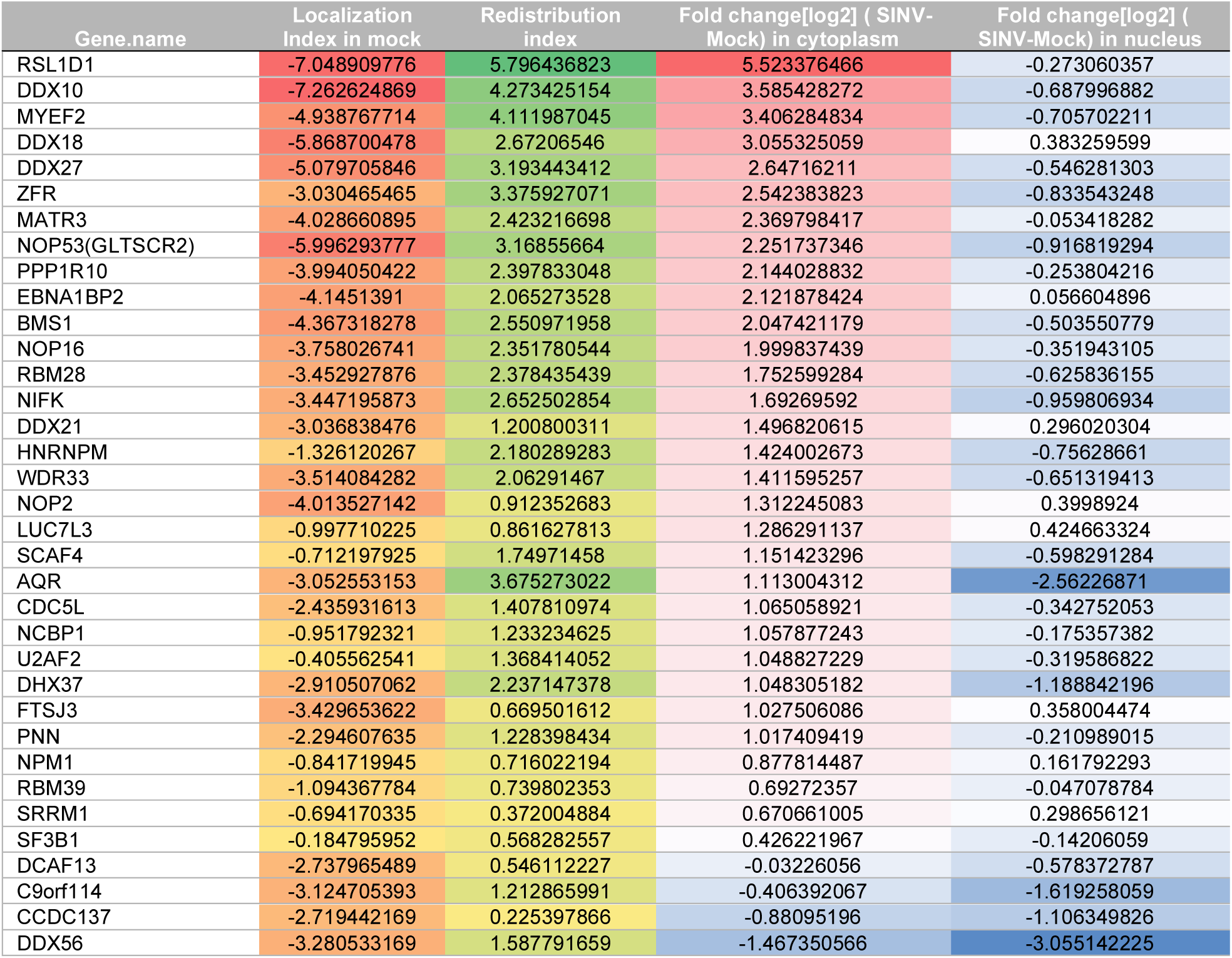
The complement of rnRBPs. This include nuclear proteins that interact with SINV RNA and are redistributed to the cytoplasm in SINV-infected cells. Localisation index (cytoplasmic/nuclear protein intensity ratio in mock cells), redistribution index (cytoplasmic/nuclear protein intensity ratio in infected versus mock cells)

To validate our proteomic results, we followed two orthogonal approaches. First, we fractionated nucleus and cytoplasm and detected by western blotting 7 rnRBPs (MATR3, AQR, SF3B1, NOP2, NOP53, NIFK, DDX21) and a cytoplasmic (GAPDH), a nuclear envelope (LamininB), a nucleolar (FBL) and a nucleoplasmic (ZC3H11A) control. All the 7 rnRBPs displayed an increased presence in the cytoplasm after SINV infection, while the controls showed no changes in localisation (Figure 4I). Second, we infected cells with SINV_nsP3-mScarlet_ and detected the location of three SINV RNA interactors: two nuclear (NOP53 and NOP2), and one cytoplasmic (PURB); adding the nucleoplasmic RBP ZC3H11A as a negative control. While NOP53 and NOP2 were prominently nucleolar in uninfected cells, both proteins exhibited a cytoplasmic population in infected cells (Figure 5A-B). Interestingly, cytoplasmic NOP53 and NOP2 fully co-localised with nsP3-mScarlet, further confirming the engagement of rnRBPs with the RdRp complex and the ROs. PURB was prominently cytosolic in uninfected cells, and it redistributed to the ROs upon infection co-localising with nsP3-mScarlet (Figure 5C). No changes in localisation were observed for ZC3H11A, which neither interact with SINV RNA nor nsP3-mScarlet (Figure 5D).

**Figure 5.**
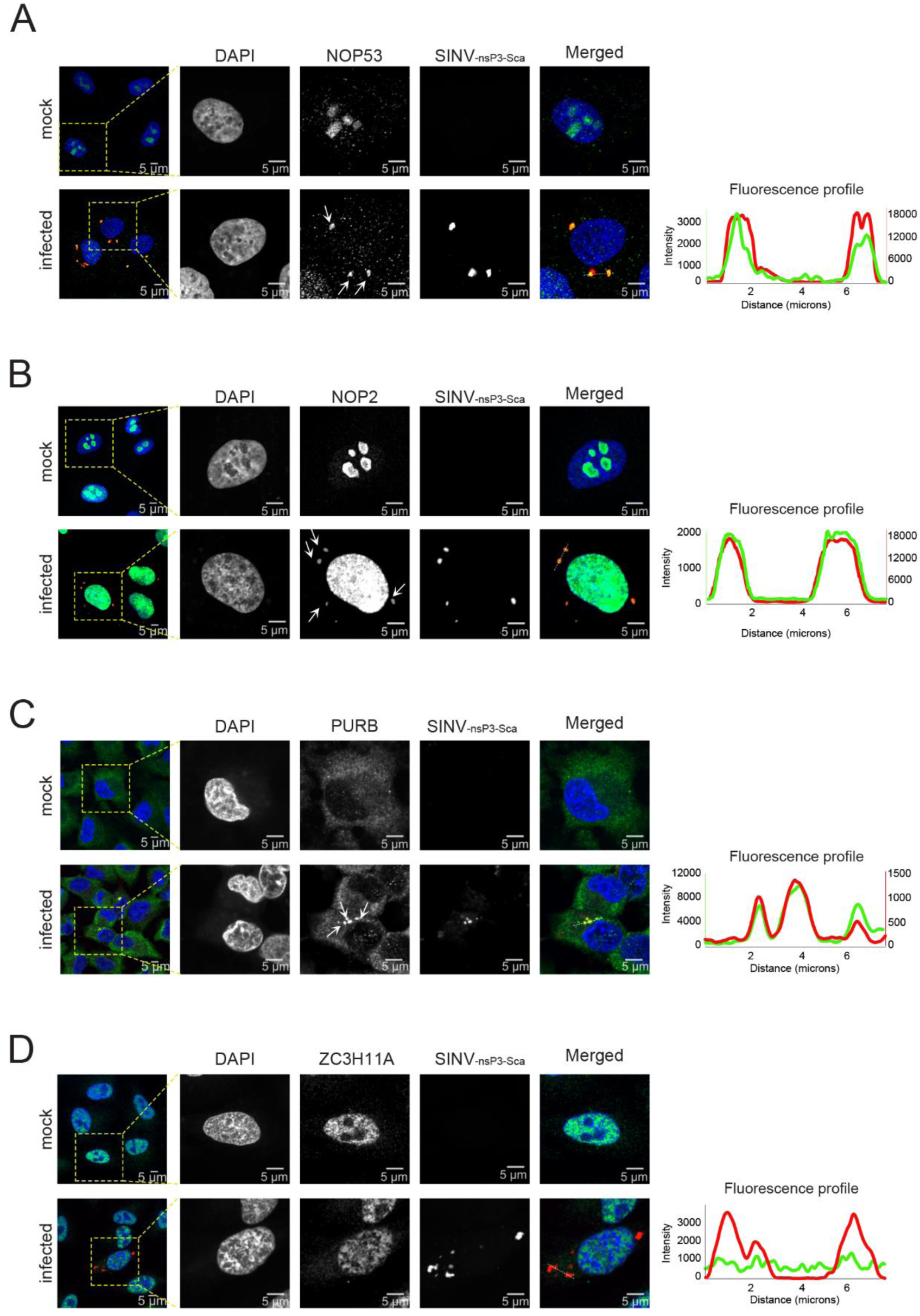
Analysis of the subcellular location of SINV RNP components by immunofluorescence. A549 cells were infected with SINV_nsP3-mScarlet_ (MOI= 3) and at 24hpi were processed for immunofluorescence using nsP3-mScarlet (nsP3-Sca) to localise the ROs and DAPI to delimit the cell nucleus. A and B) analysis of the nucleolar rnRBPs NOP53 and NOP2. C) Analysis of the cytosolic SINV RNA-binder PURB. D) Negative control with a nuclear ZC3H11A that does not interact with SINV RNA.

Altogether, our data reveal that a subset of nuclear RBPs is specifically relocated to the cytoplasm in infected cells, where they interact with vRNA and/or the RdRp complex. The fact that most of the RBPs translocated to the cytoplasm interact with vRNA or/and RdRp complex suggest that they may be recruited and retained by these viral molecules.

### SINV vRNP components display broad spectrum regulatory activities

To determine if the components of SINV RNPs are important for infection, we performed a metanalysis of 73 genome-wide screenings of viral fitness that used different methodologies (i.e., siRNA, CRISPR/cas9) and viruses (36 viruses from 18 families; Figure S5A-B). There is a limited overlap between these datasets when compared one-to-one, even when datasets for the same virus are used (Figure S5C). This is probably due to the genome-wide screenings being prone to high incidence of false positives and negatives. Nevertheless, we found that RBPs within vRNPs produced far more phenotypes across datasets than their cRNP counterparts (Figure 6A). Because *bona fide* regulators of virus infection are expected to be present in several independent screens, we classified as ‘potential regulators’ the RBPs whose involvement in infection is supported by at least three genome-wide screens, while considering ‘master regulators’ those with phenotypes spanning three or more viral families. Importantly, SINV RNP components were significantly enriched in both categories when compared to the cRNPs (Figure 6B and S5D), linking viral fitness phenotypes to vRNA binding. To test specifically if translocation of nuclear RBPs to the cytoplasm is important for SINV, we performed siRNA knockdown (KD) experiments targeting six rnRBPs (NOP2, NOP53, AQR, SF3B1, MATR3 and DDX21), one cytosolic RBP (MKRN2) and a nuclear RBP that neither bind to SINV RNA nor relocate to the cytoplasm (ZC3H11A). Transient siRNA KD of selected targets did not affect cell viability (Figure S5F). Depletion of any of the six rnRBPs caused a consistent and robust increase in viral proteins and RNA as well as virus titre (Figure 6B-I). Conversely, KD of the cytosolic MKRN2 hindered SINV gene expression (Figure S5G), while the negative control ZC3H11A had no effect as previously shown for SFV (Figure S5H) (Younis et al., 2018). These results suggest that rnRBPs possess moonlighting antiviral activities.

**Figure 6.**
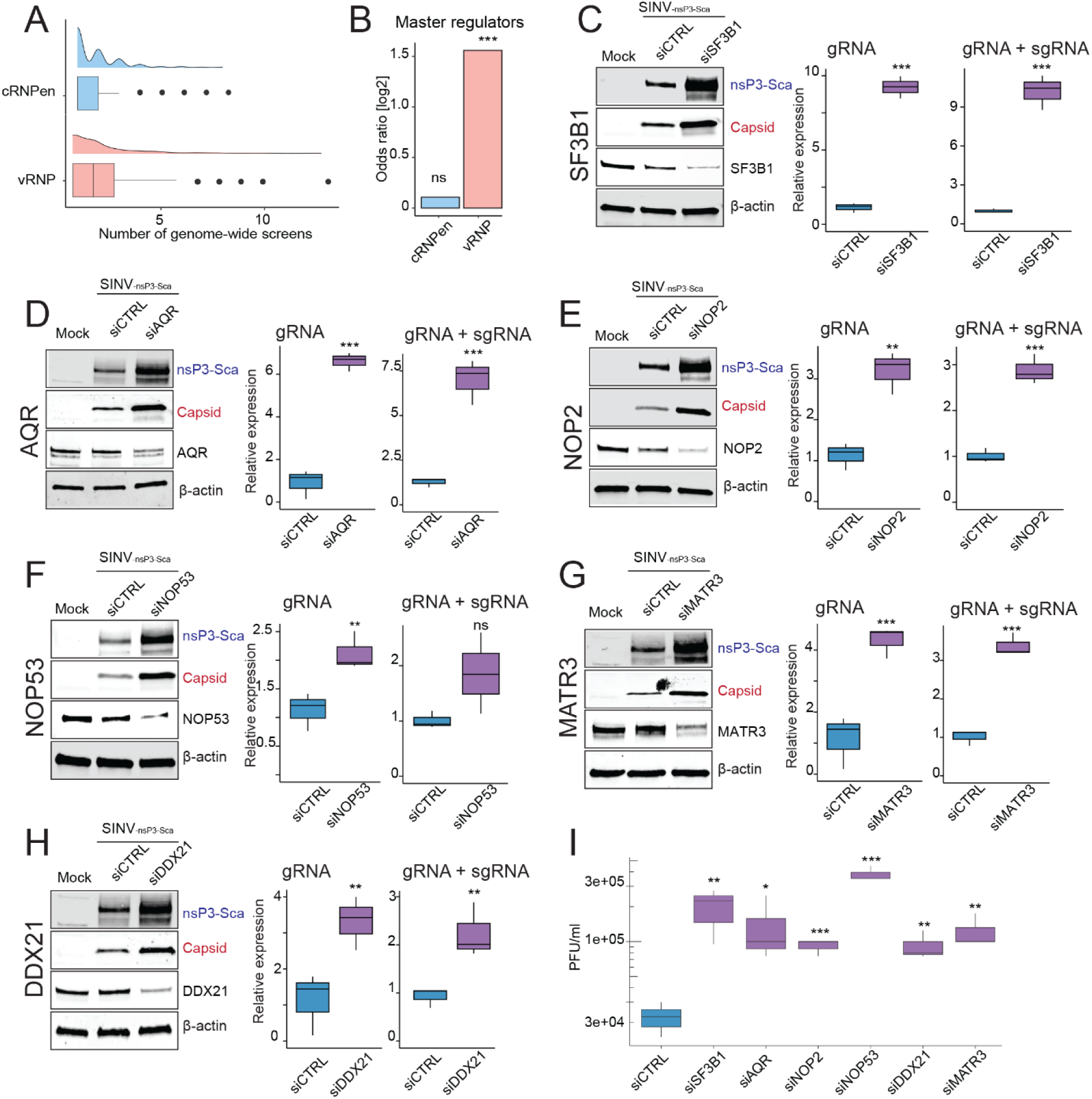
Redistributed nuclear proteins suppress SINV infection. A) Distribution of the frequency of genome-wide screens reported “hits” per each protein within vRNP or cRNPen datasets. B) Bar plot showing the enrichment of master regulators (proteins linked to three different virus families) in vRNP or cRNPen datasets. Ns and *** indicate non-significant, p-value <0.01 respectively estimated by Welch’s t-test C–H) HEK293 cells were treated with non-targeting siRNA (siCTRL) or with an siRNA against protein of interest (siPOI) for 24 hours, followed by infection with SINV_nsP3-mScarlet_ (MOI = 0.5) for 18 hpi. Left panels show immunoblotting analysis. Right panels show qRT-PCR analysis with primers against SINV genomic (gRNA) or genomic and sub-genomic (gRNA + sgRNA) RNAs, n=3. *, ** and *** indicate student t-test p-value < 0.1, 0.05 and 0.01 respectively. I) Titration the viral particles from supernatants of HEK293 cells treated with the different siRNAs and infected with SINV_nsP3-mScarlet_ (MOI= 0.5) for 18 hpi, n=3. *, ** and *** indicate student t-test p-value < 0.1, 0.05 and 0.01 respectively. Related to Figure S5.

We next expanded the functional characterisation of the rnRBPs to other alphaviruses (SFV and CHIKV), the picornavirus coxsackievirus B3 (CVB3) and adenovirus serotype 5 (HAdV-5). SFV, CHIKV and CVB3 are cytoplasmic, positive-sense single-stranded (ss)RNA viruses as SINV, while HAdV-5 is a nuclear double-stranded (ds)DNA virus. We found that SF3B1, AQR, MATR3 and NOP2 KD caused increased SFV and CVB3 gene expression, as occurred with SINV (Figure 7A, B and C). However, only NOP2 KD and, to a lesser extent, MATR3 caused a substantial increase in CHIKV capsid protein production (Figure 7D). rnRBP KD had no effect on the HAdV-5 with the exception of SF3B1 that is required for vRNA splicing (Figure S6A) (Westergren Jakobsson et al., 2021). Our data thus reveal that the antiviral activities of rnRBPs span RNA virus species and families.

**Figure 7.**
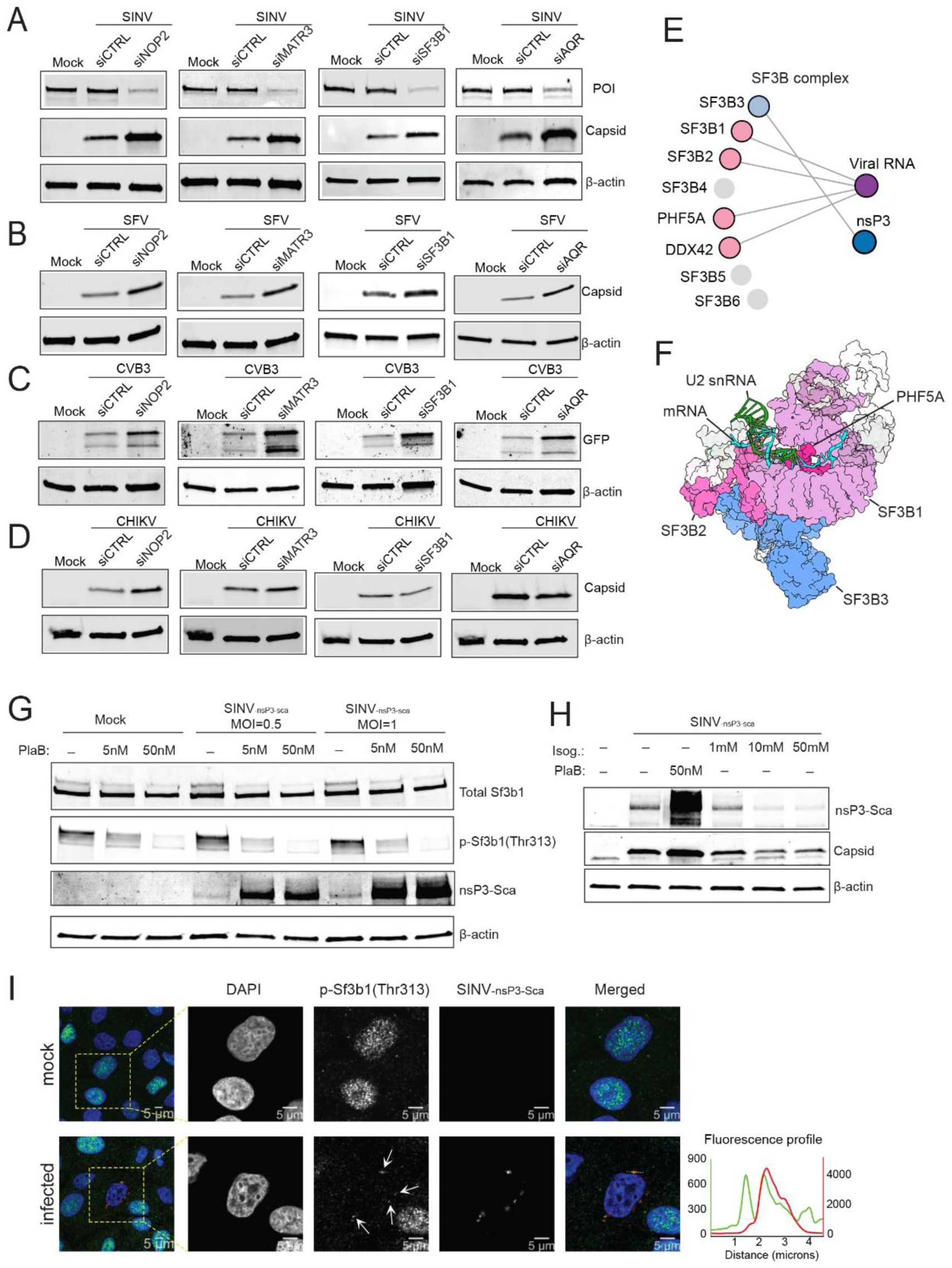
Components of SINV RNPs as pan-viral regulators. A-D) HEK293 cells treated with siRNA against nontargeting (siCTRL) or protein of interest (siPOI) for 48 hours, followed by infection with SINV, SFV, CVB3 and CHIKV (MOI= 0.1, 18hpi). E) Schematic of SF3B complex components (CORUM database id: 1737), highlighting detected interactions with SINV RNA or nsP3. F) Crystal structure of human pre-Bact-2 spliceosome showing selected components SF3b complex with U2 snRNA and mRNA; PDB ID: 7ABH. G)Immunoblotting analysis of total SF3B1, p-SF3B1(Thr313), nsP3-Scarlet (nsP3-Sca) and β-actin from mock or SINV_nsP3-mScarlet_ (MOI = 0.5 and 1) infected A549 cells, pre-treated for 2 hours with DMSO or Pladienolide B (PlaB) at 5 and 50 nM. H)Immunoblotting analysis of nsP3-Scarlet (nsP3-Sca), capsid and β-actin from mock or SINV_nsP3-mScarlet_ (MOI= 1) infected A549 cells, pre-treated for 2 hours with DMSO or Pladienolide B (PlaB) or Isoginkgetin (Isog.) at the indicated concentrations. I) A549 cells were infected with SINV_nsP3-mScarlet_ (MOI= 3) and at 24hpi were processed for immunofluorescence using p-SF3B1(Thr313) antibody, nsP3-mScarlet (nsP3-Sca) to localise the ROs, and DAPI to delimit the cell nucleus. Related to Figure S6.

### The SF3B complex inhibits viral gene expression in a splicing-independent manner

SF3B1 assembles with other RBPs to form the SF3B complex, which is critical in the formation of the lariat at the branching point during pre-mRNA splicing (Sun, 2020). Four subunits of this complex were detected in the SINV RNPs (SF3B1, SF3B2, PHF5A and DDX42), and a fifth interacts with nsP3 (SF3B3) (Figure 7E-F). Biochemical and structural work revealed that SF3B1 and PHF5A form a cavity that bind the intronic sequence, and the compound Pladienolide (Pla)B can compete for this cavity and inhibit splicing (Figure S6B) (Effenberger et al., 2017). To determine if SF3B complex has antiviral activity using an orthogonal approach, we treated cells with PlaB and infected with SINV. PlaB treatment caused dephosphorylation of SF3B1, which is linked to deficient splicing activity (Figure 7G) (Girard et al., 2012; Wang et al., 1998). SF3B inhibition correlated with a strong increase SINV nsP3 production (Figure 7G), phenocopying SF3B1 KD. The enhancement of SINV infection by PlaB was reproduced in two different cell lines (Figure 7G and S6C).

To assess if SINV enhancement is caused by dysregulation of splicing or a direct SF3B function on vRNA, we used isoginketin, which inhibits splicing at a downstream step (stalls spliceosome assembly at complex A, before recruitment of the U4-6 tri-snRNP) (O’Brien et al., 2008). Interestingly, isoginketin treatment did not enhance SINV gene expression and, at a high dose even repressed it, indicating that the inhibition of splicing per se does not cause the antiviral phenotype (Figure 7H). Additionally, we compared the distribution of the catalytically active phosphorylated form of SF3b1, pT313-SF3b1, during mock and SINV-infection. pT313-SF3b1 was mostly nuclear in uninfected cells, accumulating in nuclear granules compatible with speckles (Figure 7I). Strikingly, pT313-SF3b1 signal became more cytoplasmic in SINV infected cells, accumulating in the ROs. This suggests the presence of active SF3B in the ROs bound to SINV RNAs. Altogether, our data reveal a novel antiviral activity for the SF3B complex that appears to be independent of nuclear RNA splicing.

## Discussion

Recent data highlighted vRNA as a hub for critical host-virus interactions. We applied a systematic and comprehensive approach to elucidate the composition of the SINV RNPs using vRIC. In parallel, we also identified the RNA polymerase II- dependent (cRNP) and independent (mitoRNP) cellular RBP subpopulations, the latter is normally underrepresented in standard RIC studies. Our data revealed that SINV RNAs engage with over four hundred cellular RBPs, many of which also interact with the viral RdRp complex. SINV RNPs partially overlap with cRNPs, but also show important differences.

Several reports have proposed a non-canonical cap-dependent mechanism for alphaviruses (Carrasco et al., 2018), however, the components that associate with the translating SINV RNAs have not be assessed biochemically. Here, we revealed that several core EIFs, such as the cap-binding protein EIF4E, the helicase EIF4A2 and the Met-tRNAi depositing complex EIF2S1 (EIF2α) and EIFS2 (EIF2β) are not detected in SINV RNPs, while present in cRNPs. The lack of these EIFs in SINV RNPs explains why SINV sgRNA translation does not rely on EIF4E (Castello et al., 2006), EIF4A (Garcia-Moreno et al., 2013) and EIF2α (Ventoso et al., 2006). SARS-CoV-2 RNPs also lack EIF4E, suggesting that non-canonical cap-dependent mechanisms could apply to a wide range of RNA viruses (Flynn et al., 2021; Kamel et al., 2021; Lee et al., 2021; Schmidt et al., 2021). Interestingly we showed that EIF3D is upregulated during SINV infection (Garcia-Moreno et al., 2019) and here we show it is present in SINV RNPs. EIF3D can mediate cap-dependent translation (Lee et al., 2016), and whether it initiates translation for capped viral RNAs as proposed for human cytomegalovirus transcripts (Thompson et al., 2022) deserves further consideration .

Furthermore, we noticed a striking increase in the incidence of PTM enzymes in SINV RNPs, particularly kinases. The key innate immunity kinase EIF2AK2 (PKR) is activated by binding to dsRNA, which is produced as intermediate products of viral replication (Taylor et al., 2005). Our data show interaction of EIF2AK with SINV RNAs, which can explain the strong phosphorylation of EIF2S1 (EIF2α) found in SINV-infected cells (Ventoso et al., 2006). SRPK1 phosphorylates SR repeats present in many splicing factors (SRFs). It locates to SINV ROs and promotes SINV gene expression (Garcia-Moreno et al., 2019). This study now shows that SRPK1 directly engages with SINV RNAs, potentially regulating the microenvironment of the ROs. Other kinases with no characterised RNA-binding activity were identified in SINV RNPs, including mitogen-activated protein kinases (MAP4K4, TNIK, TAOK2 and PAK4) that regulate inflammatory responses (Kyriakis and Avruch, 2012), and cyclin-dependent kinases (CDK1, 9, 12 and 13) that control the cell cycle (Gutierrez-Chamorro et al., 2021). Whether these kinases are sequestered in the ROs to interfere with inflammation and cell cycle should be further investigated.

Unexpectantly, we found that several nuclear RBPs (rnRBPs) interact with SINV RNAs in the cytoplasm. Nuclear/cytoplasmic relocation has been reported for a few individual proteins (Lloyd, 2015; Sanz et al., 2015). To our knowledge, no systematic and comprehensive study of this phenomenon has been attempted until now. Our nuclear/cytoplasmic fractionation experiment substantially extended the range of RBPs that become cytoplasmic in SINV infected cells. We also show that most of the rnRBPs interact with SINV RNA and/or the RdRp complex, and for NOP53 and NOP2 we observed an accumulation at the ROs by microscopy. This suggests that viral RNA and proteins may recruit and retain rnRBPs in the cytoplasm. Depletion of rnRBPs strongly enhance SINV infection, evoking moonlighting antiviral roles. These virus-repressive activities span RNA virus species and families, suggesting fundamental although poorly understood antiviral mechanisms. Given the variety of rnRBPs, it is plausible that several distinct pathways operate simultaneously to repress infection. Amongst the rnRBPs we noticed an enrichment of two main protein groups: splicing and ribosomal biogenesis factors (nucleolar proteins). The nucleolus has been related to stress sensing and, in some instances, it involves the translocation of nucleolar proteins such as NOP53 to the cytoplasm (Lee et al., 2012; Lin et al., 2022; Yang et al., 2016). The accumulation of NOP53 at the ROs suggests that such sensing mechanism may also operate in virus infected cells and could involve vRNA.

Our study also revealed a set of splicing factors that engage with SINV RNAs, with a highlight for the SF3B complex with several components identified. SF3B is a central component of the U2 small nuclear (sn)RNP that is required for the formation of the lariat at the branching point (Sun, 2020). KD of the central component of the SF3B complex SF3B1 causes a strong enhancement of SINV virus gene expression, implying that it displays an antiviral role. This is confirmed orthogonally by treatment with PlaB inhibitor, indicating that the association of SF3B with vRNA is important for the antiviral phenotype. SF3B effects in infection could be direct by acting on vRNA, or indirect by a dysregulation of splicing. SINV causes a potent inhibition of cellular transcription, suggesting that splicing regulation might not be relevant for the phenotype (Gorchakov et al., 2005). Moreover, treatment with isoginketin, which blocks splicing at a downstream step does not phenocopy SF3B1 KD, suggesting that dysregulation of splicing does not suffice to enhance SINV gene expression. The function of the SF3B complex on SINV RNAs and whether it requires the U2 snRNA remains to be elucidated.

## ACKNOWLEDGEMENTS

A.C. is funded by the European Research Council (ERC) Consolidator Grant ‘vRNP-capture’ 101001634 and the MRC grants MR/R021562/1 and MC_UU_00034/2. W.K. is funded by the European Union’s Horizon 2020 research and innovation programme under Marie-Sklodowska-Curie n# 842067. V.R. is funded by the European Union’s Horizon 2020 research and innovation programme under Marie-Sklodowska-Curie n#892756. A.E.B. is funded by Fundación Ramón Areces post-doctoral fellowship programme. L.I. is funded by the BBSRC DTP scholarship number BB/M011224/1. I.D. is funded by the Wellcome Investigator Award 209412/Z/17/Z. We thank F. J. M. Van Kuppeveld for the CVB3-eGFP construct. We thank Claudia Guida for her invaluable feedback in the elaboration of the manuscript.S.M.2, Y.D. and E.K .are funded by an EPSRC grant (V011359/1 (P).

## AUTHOR CONTRIBUTIONS

Conceptualization, W.K., S.M.^2^, A.C.; Methodology, W.K., M.G.M., V.R., A.I.J. S.M.^2^, I.D, A.C.; Investigation, W.K., V.R., A.E.B., Z.R.L., Y.D., M.G.M., M.M, L.I., S.M.^1^, A.I.J., M.H., E.K., S.M.^2^, A.C.; Writing, original draft, W.K. and A.C.; Writing, editing, W.K., V.R. A.E.B., Z.R.L., Y.D., M.G.M., A.M., S.M., S.M.^2^, A.C.; Funding acquisition, W.K., S.M.^2^, A.C.; Resources, A.M., I.D., S.M.^2^, A.C.; Supervision, W.K., S.M.^2^, A.C.

## MATERIALS AND METHODS

### Experimental model and subject detail

#### Cell culture

HEK293 (Flp-In T-REx, Thermo Fisher Scientific # R78007) and A549 cells were maintained in DMEM (Gibco, 41965039) with 10% fetal bovine serum (FBS) (Gibco, 10500064) and 1x penicillin/streptomycin (Sigma Aldrich, P4458) at 37**°**C with 5% CO_2_. BHK21 and HEK293T cells, for growing virus stocks, were maintained in same conditions.

#### Viruses

Alphaviruses used were: Sindbis virus (SINV) that was produced by in vitro transcription from the plasmid pT7svwt (Castello et al., 2006); chikungunya virus (CHIKV) belongs to the Eastern/Central/South African (ECSA) genotype, ICRES-1 strain. Replicative alphaviruses with fluorescent reporters (SINV_mCherry_ and SFV_mCherry_) were generated by duplication of subgenomic promoter and insertion of mCherry (Garcia-Moreno et al., 2019). The SINV_nsP3-mScarlet_ was generated by amplifying mScarlet sequence from pUC19_CRISPR_CT_Scarlet_GSG_P2A_BSD (primer forward: GGGGACTAGTATGGTGAGCAAGG; reverse: ccccACTAGTCTTGTACAGCTCG) and cloned into pT7SINV-WT using SpeI restriction enzyme. Coxsackie B3 virus (CVB3) has eGFP inserted at the N-terminus followed by a 3Cpro cleavage site at N-terminus (Lanke et al., 2009). For adenovirus infections, adenovirus serotype 5, HAdV-5, which belongs to subgroup C was used.

### Viral RNA interactome capture

HEK293 (FLP-IN T-REX) cells were grown in sets of 4x15 cm dishes. Cells were infected with SINV for 1h with a multiplicity of infection (MOI=1). After 2 hours, supernatant was replaced with fresh media supplemented with 20µM flavopiridol hydrochloride hydrate (Fvo, Cat.No. F3055, Sigma-Aldrich) and incubated at 37°C for 3 hours. Then the supernatant was replaced with fresh media supplemented with 20µM Flavopiridol hydrochloride hydrate (Fvo, Cat.No. F3055, Sigma-Aldrich) and 100µM 4-Thiouridine (4SU, Cat.No. T4509, Sigma-Aldrich), an incubated at 37°C. At 24hpi, media was discarded, and cells were rinsed once with PBS (Phosphate-buffered saline). After removal of the PBS, the cell monolayer was irradiated twice with 200 mJ/cm2 using ultraviolet light at 365nm. At this stage, cells were lysed with 8mL of lysis buffer (20mM Tris-HCl pH 7.5, 500mM LiCl, 0.5% LiDS wt/vol, 1mM EDTA, 0.1% IGEPAL (NP-40) and 5mM DTT), homogenised by passing several times through a 5 mL syringe equipped with 27G needle, and polyadenylated RNA purified with oligo(dT) magnetic beads as in (Castello et al., 2013; Garcia-Moreno et al., 2019). Then homogenized lysate incubated with 0.8mL of pre-equilibrated oligo(dT)25 magnetic beads (New England Biolabs, #S1419S) for 1h at 4°C with gentle rotation. Beads were collected by magnet and the lysate was transferred to a new tube and stored at 4°C. Beads were washed for 5min at 4°C with gentle rotation once with 5mL of lysis buffer, followed by two washes with 5mL of buffer 1 (20mM Tris-HCl pH 7.5, 500mM LiCl, 0.1% LiDS wt/vol, 1mM EDTA, 0.1% IGEPAL and 5mM DTT), and two washes with buffer 2 (20mM Tris-HCl pH 7.5, 500mM LiCl, 1mM EDTA, 0.01% IGEPAL and 5mM DTT). Beads were then washed twice with 5 mL of buffer 3 (20mM Tris-HCl pH 7.5, 200mM LiCl, 1mM EDTA and 5mM DTT) at room temperature for 3min. Beads were resuspended in 300μL of elution buffer (20mM Tris-HCl (pH 7.5) and 1mM EDTA.) and incubated for 3min at 55°C with agitation. After collecting the beads with a magnet, supernatants with the eluted proteins were collected and stored at −80°C. The lysates were subjected to a second round of capture repeating the same protocol and the eluates from the first and second capture cycles were combined. Prior to mass spectrometry sample processing, samples were RNase treated with ∼0.02U RNase A and RNase T1 at 37°C for 1h. The controls described in Figure 1B were generated as above but omitting the virus, Fvo or/and UV irradiation.

### Subcellular fractionation

HEK293 (FLP-IN T-REX) cells (4X10^6^) were infected with SINV (MOI=10) or mock-infected for 24 hours. To separate the cytoplasmic and nuclear fractions, cells were harvested and suspended in 1X PBS, followed by centrifugation at 100g for 5 min at 4^ο^C, the cell pellet was incubated with 500 µL lysis buffer (1% IGEPAL, 150mM NaCl, 50mM HEPES pH 7.9, 0.1mM AEBSF serine protease inhibitor) at 4^ο^C for 10 mins. Followed by centrifugation at 7000g for 5min, the cytoplasmic supernatant (cytoplasmic fraction) was transferred to new tube and cleared by centrifugation at 15,000xg at 4^ο^C for 5 min. The pellet (nuclear fraction) was suspended in 500µL RIPA buffer (0.1%SDS, 0.5% sodium deoxycholate, 150mM NaCl, and 50mM HEPES, pH 7.9 and 0.1mM AEBSF serine protease inhibitor) supplemented with Benzonase (1U/mL) and incubated on ice for 1hr then supplemented with SDS and DTT to final concentration of 1%SDS and 0.1M DTT. Next, Nuclear lysate was boiled for 2min and cleared by centrifugation at 15,000g for 5min. For proteomic analysis four biological replicates were generated.

### SINV-nsP3 mScarelt-tagged immunoprecipitation

1×10^7^ cells HEK293 FLP-IN T-REX cells were infected with SINV_nsP3-mScarlet_ (bait) or SINV_wt_ (negative control) at 1 MOI. After 18 hours, cells were scrapped and spin down at 300 g 4°C for 10 mins. After the centrifugation, supernatants were discarded and cell pellet washed once with 1X PBS, followed by another centrifugation at 300xg 4°C for 10min. After discarding the supernatant, cell pellets were lysed with 200µL ice-cold lysis buffer (10mM Tris-Cl pH 7.5, 150mM NaCl, 0.5mM EDTA, 0.5% IGEPAL, benzonase(1U/mL) and 0.1mM AEBSF serine protease inhibitor), and incubated on ice for 30min. Then, cell lysates were cleared by centrifugation 17,000xg for 10min at 4°C. The supernatants were transferred to new tubes and diluted 1:1 with the dilution buffer (10mM Tris-Cl pH 7.5 150mM NaCl 0.5mM EDTA 0.05% IGEPAL). Next, cell lysates were incubated with 15µL of RFP-Trap® Magnetic Particles M-270 (pre-washed once with the dilution buffer) for 1 hour at 4 °C with mild rotation. After the incubation, the magnetic beads were washed five times with wash buffer1 (10mM Tris/Cl pH 7.5, 150mM NaCl, 0.5mM EDTA and 0.25% IGEPAL), followed by two washes with the dilution buffer. The proteins elution from the beads was done twice at room temperature with shaking, the first elution with 45µL 0.2M glycine pH 2 solution and the second elution with 45µL 0.2M glycine pH 2 (supplemented with 1%SDS) solution. Finally combined protein eluates were neutralized with 10µL 1M Tris-base pH 10. For proteomic analysis three biological replicates were generated.

### siRNA-transfection

HEK293 FLP-IN T-REX cells were seeded in 24-well plate, 2X10^5^ cells per well. 24h later, cell media is exchanged with fresh media (supplemented with 5% FBS and 1x penicillin/streptomycin). After one hour, cells were transfected with siRNAs (Table S8), at final 50 nM final concentration, using X-tremeGENE™ 360 transfection reagent according to the manufacturer instructions and incubated at 37**°**C with 5% CO_2_ for the indicated time periods. At the time of infection, the FCS concentration was further reduced to 2.5% and incubated at 37**°**C with 5% CO_2_ for 18h.

### Plaque assay

Vero cells were seeded on 12 well plates (2×10^5^ cells/well). Next day a 10-fold dilution of the infectious supernatant was performed, and 400uL used to infect each well. Later cells were incubated for 1h at 37°C and 5% CO_2_, rocking the plate every 15 minutes. After infection, 1mL overlay of 0.6% Avicel immobilization media (1:1:2 ratio of 2.4% room temperature Avicel stock, 1X PBS and warmed 2x MEM media supplemented with 4% FBS), was added and the plates incubated for a further 48 hours at 37°C and 5% CO_2_. The immobilizing media was aspirated and discarded, and cells fixed with 8% formaldehyde for 1h. The fixing agent was then aspirated and discarded appropriately before staining with crystal violet solutions for at 15 min. The crystal violet stain was then washed off and the plates left to dry before counting of the plaques.

### Cell lysis and Western blot

To prepare whole cell lysate for western blotting, virus- and mock-infected cells were harvested at the times indicated as indicated in the figure legeneds, washed once with 1X PBS and resuspended in RIPA buffer (0.1%SDS, 0.5% sodium deoxycholate, 150mM NaCl, and 50mM HEPES, pH 7.9 and 0.1mM AEBSF serine protease inhibitor) supplemented with Benzonase (1U/mL) and incubated on ice for 1h, then supplemented with SDS and DTT to final concentration of 1%SDS and 0.1M DTT. Next, total lysate was boiled for 2min and cleared by centrifugation at 15,000xg for 5min. Western blotting was performed as previously described in (Garcia-Moreno et al., 2019), antibodies used are listed in Table S8.

### Immunofluorescence and single molecule in situ hybridisation

Immunofluorescence was performed as previously described (Garcia-Moreno et al., 2019), with minor modifications. 1.5×10^5^ A549 cells were grown on sterile 12mM, thickness 1.5H round-glass coverslips (Marienfeld, #0117520) in a 24-well plate. Cells were infected with SINV _nsP3-mScarlet_ (MOI= 1). Cells were fixed 18 hpi in 4% paraformaldehyde for 15 min at room temperature and permeabilised with Digitonin (Sigma Aldrich) (1.25 µg/ml) for 10 min at room temperature. Cells were washed twice in 1X PBS and then incubated in 5% BSA in PBS for 30 min at room temperature. Cells were incubated for 1 h at room temperature with primary antibody solution (5% BSA, 0.1% Tween 20, in PBS). Primary antibodies used are listed in Table S8. Cells were then washed three times with 0.1% Tween-20 in PBS and then incubated for 1 h with the appropriate AlexaFluor-fluorescently conjugated secondary antibody in (5% BSA, 0.1% Tween 20, in PBS) (Table S8) (from now on cells were kept in the dark). Cells were then washed three times with 0.1% Tween-20 in PBS and once in 1X PBS. Cells were incubated with DAPI (1mg/ml) for 10 min at room temperature, washed once in 1X PBS and dipped in water. Coverslips were mounted using ProLong Diamond antifade mounting media (Invitrogen).

smFISH was performed as previously described (Lee et al., 2022), with minor modifications. 1.5×10^5^ A549 cells were grown on sterile 12mM, thickness 1.5H round-glass coverslips (Marienfeld, #0117520) in a 24-well plate. Cells were infected with 1 MOI SINV_nsP3-mScarlet_. Cells were fixed 18 hpi in 4% paraformaldehyde for 15 min at room temperature and permeabilised in 0.1% Triton X-100 in PBS for 10 min at room temperature. Cells were washed twice in 1X PBS and twice in 2X SSC. Cells were pre-hybridised twice for 20 min at 37°C in wash solution (2× SSC, 10% formamide, pre-warmed). Hybridisation was performed using hybridisation solution (2× SSC, 10% formamide, 10% dextran sulphate) supplemented with 125 nM SINV RNA-specific Stellaris probes (LGC Biosearch Technologies) and incubated overnight at 37°C in a humidified chamber (from now on cells were kept in the dark). Cells were then washed in warm wash solution for 20 min at 37°C and incubated with DAPI (1 µg/ml) in wash solution at room temperature for 10 min. Finally, cells were washed once with wash solution for 20 min at room temperature and twice with 2× SSC for 10 min each at room temperature. Coverslips were dipped in water before mounting using ProLong Diamond antifade mounting media (Invitrogen).

### RNA extraction and qRT-PCR

Mock or virus infected cells were harvested at times indicated in figure legends, and centrifuged for 5 min at 300xg at 4**°**C. Cell pellets were washed once with PBS and RNA extraction was done using RNeasy Plus kit (Qiagen #74136) according to the manufacturer instructions. Reverse transcription and qRT-PCR analysis was performed by Luna universal one-step qRT-PCR kit (NEB # E3005L) with primers list in Table S8.

### mCherry and GFP microplate reader assay

1×10^5^ HEK293 FLP-IN T-REX GFP cells were seeded on each well of a 96-well microplate with flat mClear bottom (Greiner Bio-One, #655986) in DMEM media without phenol-red and supplemented with 10% FBS and 1mM sodium pyruvate. For mCherry- and eGFP-tagged viruses, cells are infected at MOI=1 in complete DMEM lacking phenol-red with 5% FBS. Cells were incubated at 37°C and 5% CO_2_ in a CLARIOstar fluorescence plate reader (BMG Labtech) with atmospheric control unit for the time periods indicated in the figures. For measurement of eGFP expressed from HEK293 FLP-IN T-REX cells, induction was performed with 1ug/ml doxycycline, prior to incubation in the microplate reader.

### Cell viability

Total number of cells and percentage of alive cells were estimated by trypan blue staining and Countess II FL Automated Cell Counter (Thermo Fisher Scientific).

### Mass spectrometry

Proteins samples (vRIC and nuclear cytoplasmic fractionation) were treated with benzonase (1U/mL) followed by clean-up via the bead-based single-pot, solid-phase-enhanced sample-preparation (SP3) method, using Magnetic Carboxylate Modified Particles beads (Sigma-Aldrich, cat.no.45152105050250) (Perez-Perri et al., 2021). Protein digestion was performed on beads using Trypsin Gold (MS grade; Promega, cat. no. V5280), as previously described in (Hughes et al., 2019). For processed peptides, liquid chromatography (LC) was performed using an Ultimate 3000 ultra-HPLC system (Thermo Fisher Scientific). Peptides were initially trapped in C18 PepMap100 pre-column (300 µm inner diameter x 5mM, 100A, Thermo Fisher Scientific) in Solvent A (Formic acid 0.1% (v/v), Medronic acid 5 µM). Trapped Peptides were separated on the analytical column (75 µm inner diameter x 50cm packed with ReproSil-Pur 120 C18-AQ, 1.9mM, 120 A, Dr. Maisch GmbH) in a 60 min 15%– 35% [vol/vol] acetonitrile gradient with constant 200 nL/min flow rate. Separated peptides were directly electrosprayed into a QExactive mass spectrometer (Thermo Fisher Scientific). Mass spectra were acquired in the Orbitrap (scan range 350-1500 m/z, resolution 70000, AGC target 3 x 10^6^, maximum injection time 50 ms) in a data-dependent mode. the top 10 most abundant peaks were fragmented using HCD (resolution 17500, AGC target 5 x10^4^, maximum injection time 120 ms) with first fixed mass at 180 m/z.

The nsP3 interactome proteins samples were processed similarly, but the trypsin-digested peptides were separated on Ultimate 3000 nano-LC 1000 system (thermos Fisher Scientific) equipped with a C18 PepMap100 pre-column (300 µm inner diameter x 5mM, 100A) an in-house packed C18 column (Reprosil-Gold, Dr. Maisch, 1.9 μm particle size ID: 50 μm, length: 50 cm). Mobile phase A (water and 0.1% formic acid) and mobile phase B (acetonitrile and 0.1% formic acid) were used for separation with a 15 min linear gradient of 12-38% B at a flow rate of 100 nl/min. Eluting peptides were electro-sprayed into an Orbitrap Fusion Lumos mass spectrometer (Thermo Fisher Scientific). MS1 spectra were acquired in the orbitrap (350-1400 m/z, resolution 60000, AGC target 3 x 106, maximum injection time 50 ms) in a data-dependent mode. The top 20 most abundant peaks in the survey scan were fragmented using HCD (normalized collision energy: 30%). MS2 spectra were acquired in the ion trap (scan rate: turbo, AGC target 1 x104, maximum injection time 32 ms).

### Proteomic quantitative analysis

Protein identification and quantification were performed using Andromeda search engine implemented in MaxQuant (1.6.3.4) under default parameters (Cox et al., 2011). Peptides were searched against reference Uniport datasets: human proteome (Uniprot_id: UP000005640, downloaded Nov2016) and SINV proteome derived from pT7-SVwt plasmid. False discovery rate (FDR) was set at 1% for both peptide and protein identification. MaxQuant search was performed with “match between run” activated in the searches for the subcellular fractionation experiment and nsP3 interactome data.

MaxQuant (proteinGroups) outputs were used for downstream relative quantification. Proteins flagged as potential contaminants by MaxQuant were filtered out, R-package “DEP (1.4.1)”, together with proteins with missing values across all samples. For relative quantification between Mock or infected samples, proteins raw intensities were normalized, within the same cellular fraction (in case of the fractionation experiment) and transformed using R-package Variance Stabilizing Normalization “VSN (3.50.0)”. Missing value imputation was performed only for proteins undetected in all replicates in one experimental condition, while present in the other condition (at least in 2 replicates). Imputation was implemented using local minimum determination method (Mindet) (Lazar et al., 2016). R-package “limma (3.38.3)” were used for statistical analysis for the processed protein intensities applying the empirical Bayesian method moderated t-test with p-values adjusted for multiple-testing using Benjamini-Hochberg method. For the vRIC and nsP3 interactome samples, protein intensities were processed as described above but omitting the normalization step.

### Analysis of RBP binding preferences with ENCODE eCLIP data

eCLIP bed files were downloaded from the ENCODE database use ENCODExplorer on 07/04/21. GRCh38.99 reference genome annotation from ensemble (ftp://ftp.ensembl.org/pub/release-99/gtf/homo_sapiens/Homo_sapiens.GRCh38.99.chr.gtf.gz) was used to determine the features within which binding sites occurred. This was achieved using the findOverlaps function from the genomicRanges package to align coordinates of eCLIP hits with genome annotations. To identify eCLIP hits present in introns, an intron annotation object was created by extracting the coordinates between exons of genes from the genome annotation. This was then used in the same way as the genome annotation to identify overlap with eCLIP hits. To determine the proportion of binding sites occurring within each feature, binding sites were filtered to include only those occurring in protein coding genes and counted. The frequency of binding sites in each feature was converted to a proportion of total number of binding sites in protein coding genes. ‘Binding preference’ was assigned based on the feature with the highest proportion of binding sites.

### RNA binding proteins, intrinsically disordered regions and Pfam RNA-binding domains analysis

List of RNA binding proteins identified from previous RNA interactome experiments were obtained from RBPase (Schwarzl et al., 2020), proteins detected in three independent studies were used for downstream analysis. Prediction of intrinsically disordered regions were obtained from MobiDB done by MobiDB-lite method (Piovesan et al., 2021). Assigning RNA binding proteins with either classical or non-classical domains was performed as previously described in (Kamel et al., 2021).

### PTM enzymes and protein Complexes analysis

For post-translation modification (PTM) enzymes, we focused on (de)phosphorylation and (de)ubiquitination. The list of human kinases were obtained from Kinase.com (Manning et al., 2002), phosphatases from DEPOD database (Damle and Kohn, 2019), deubiquitinases from DUP portal (Doherty et al., 2022) and Ubiquitin ligases from GO-term “0016567”. To identify cellular complexes associated with rnRBPs, a list of curated human complexes was obtained from CORUM database (Giurgiu et al., 2019).

## SUPPLEMENTARY TABLE LEGENDS

Supplementary table.1. UV-dependent RBPome. Proteins enriched in M/4SU/- over M/-/- within 1% FDR are labelled as “high confident” and 10% FDR labelled as “candidate” interactors.

Supplementary table.2. cellular (Fvo-senstive) RNPs. Proteins enriched in M/4SU/- over M/4SU/Fvo RNA interactomes within 1% FDR are labelled as “high confident” and 10% FDR labelled as “candidate” interactors.

Supplementary table.3. Mitochondrial (Fvo-insenstive) RNPs. Proteins enriched in M/4SU/ Fvo over M/4SU/- RNA interactomes within 1% FDR are labelled as “high confident” and 10% FDR labelled as “candidate” interactors.

Supplementary table.4. SINV RNPs. Proteins enriched in V/4SU/Fvo over M/4SU/Fvo RNA interactomes within 1% FDR are labelled as “high confident” and 10% FDR labelled as “candidate” interactors.

Supplementary table.5. SINV nsP3 interactome. Proteins enriched in SINV_nsP3-mScarlet_ over SINV eluates within 1% FDR are labelled as “high confident” and 10% FDR labelled as “candidate” interactors.

Supplementary table.6. Differential proteins abundance in cytoplasmic fraction during SINV infection.

Supplementary table.7. Differential proteins abundance in nuclear fraction during SINV infection.

Supplementary table.8. List of siRNA, primers, antibodies and chemical compounds used in this study.

**Figure S1.**
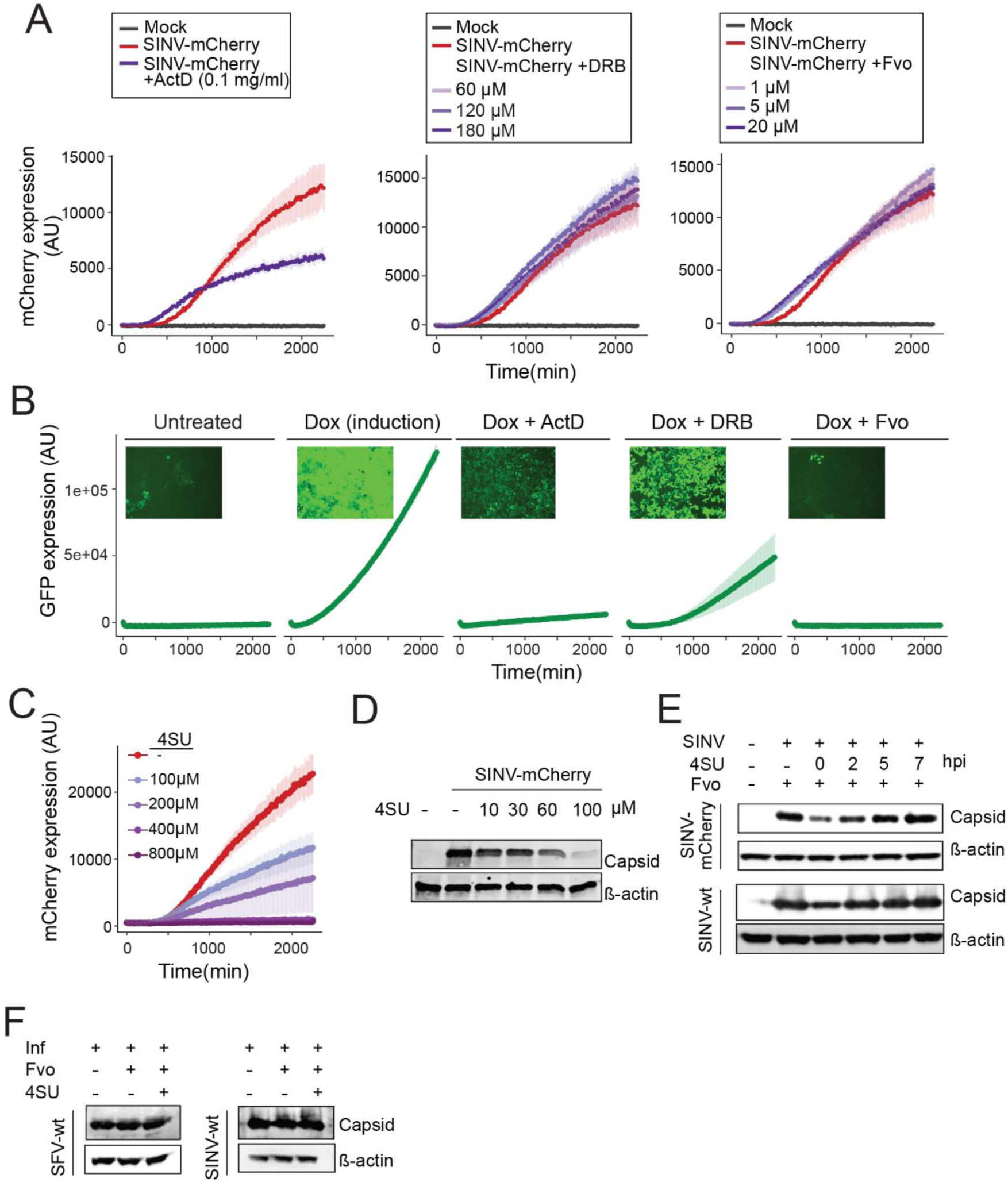
Optimisation of the vRIC experiment. A) Red fluorescent signal derived from SINV_mCherry_ infected cells treated with different RNA polymerase II inhibitors at different concentrations, Actinomycin D (ActD), 5,6-dichloro-1-β-D-ribofuranosyl benzimidazole (DRB) and Flavopiridol (Fvo). Fluorescence was measured every 15 min over a period of 36 h in a CLARIOstar plate reader with an atmospheric control unit (ACU). B) Green fluorescence derived from GFP produce after addition of deoxicycline to cultured HEK293-eGFP, in presence of the different inhibitors of RNAPs described in (A). Measurement was performed as in A. C) SINV_mCherry_ derived fluorescent signal in cells treated with different doses of 4SU (with fixed concentration of Fvo 20 µm). Measurements were performed as in (A). A-C) Error bars indicate standard deviation, n=6. D-F)Immunoblotting analysis of SINV capsid and β-actin from mock and infected cells upon treatment D) with increasing concentrations of 4SU or E) with a fix dose of 4SU added at different hpi (hour post infection). F) Immunoblot of capsid in cells infected with SINV or SFV and treated or not with Fvo and 4SU.

**Figure S2.**
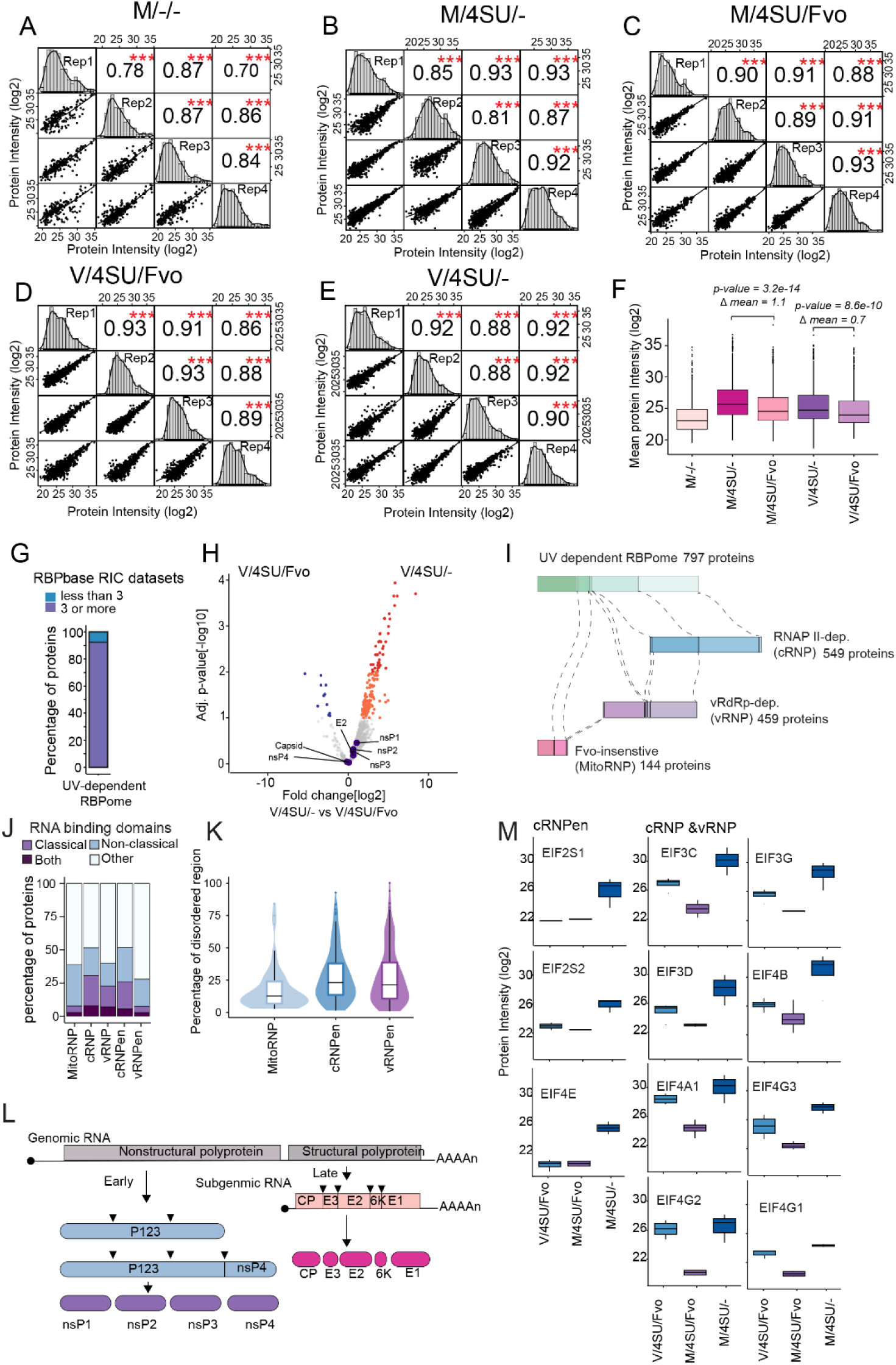
Proteomic analysis of SINV RNPs. A-E) Scatter plots showing protein intensity correlation across replicates for the different condition shown in Figure 1B. Pearson coefficient and protein intensity distribution are shown. F) Box plot showing the distribution of protein intensities in the experimental and control samples (lower and upper ends of the hinges are defined by the first and third quartile). p-values were estimated by Welch’s t-test. G) Bar plot defining the proportion of RBPs (from Figure 1F) previously reported to RBPs in RNA interactome capture (RIC) studies. H) Volcano plot showing fold change and adjusted p-value for each protein (dot) comparing V/4SU/- versus V/4SU/Fvo. Proteins enriched within 1% FDR are coloured in red and blue and those within 10% FDR in orange and cyan. Viral proteins are indicated. I) Schematic representation of different populations of RBPs identified in this study, highlighting shared proteins. J) Proportion of proteins with classical RBDs, non-classical RBDs, both or none in mitoRNPs, cRNPs, vRNPs, cRNP-enriched (en) and vRNPen. K) Prevalence of intrinsic disorder regions (IDRs) in mitoRNPs, cRNPen and vRNPen. L) Schematic of SINV gRNA, sgRNA and derived viral proteins. M) Boxplots showing protein intensities for selected eukaryotic translation initiation factors either only enriched in cellular RNPs (cRNPen) or in both viral and cellular RNPs (cRNP and vRNP).

**Figure S3.**
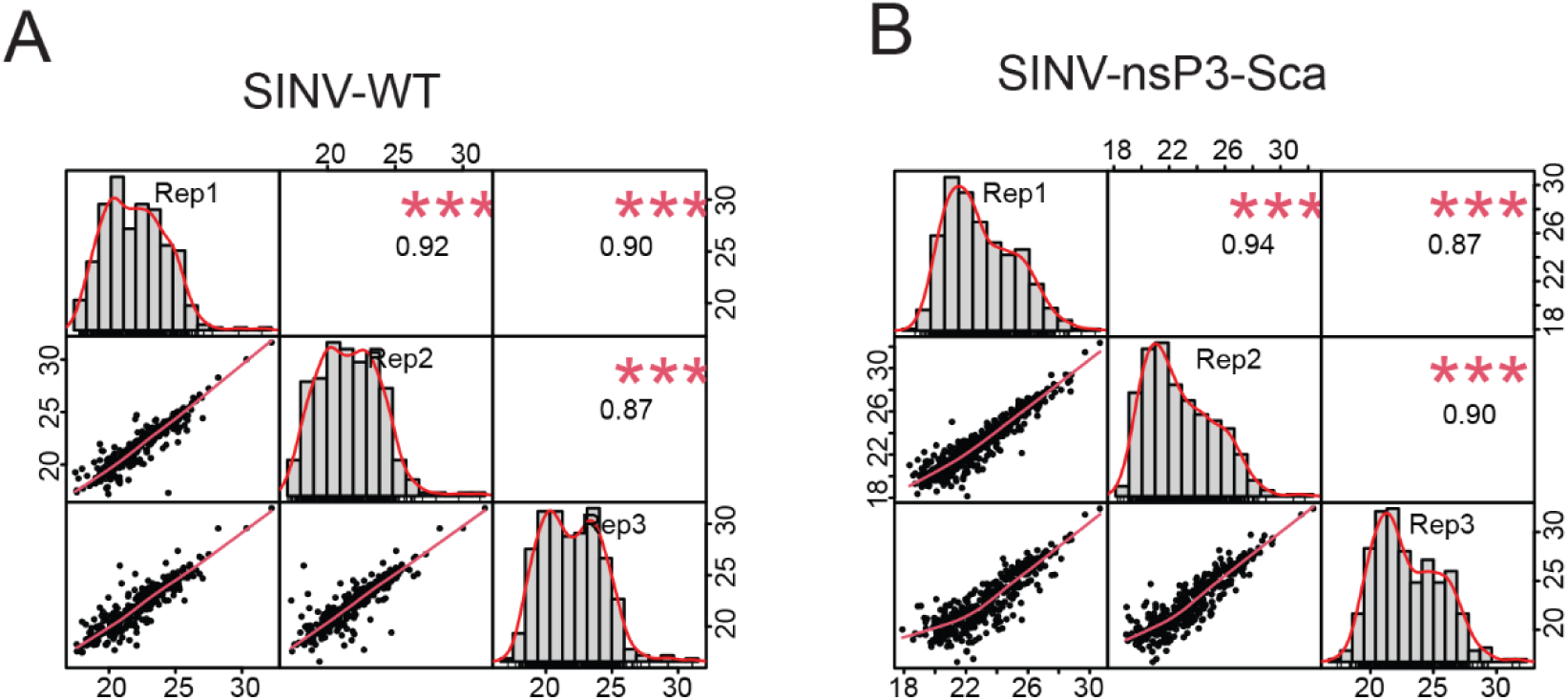
Proteomic analysis of nsP3 interacting proteins in SINV-infected cells. A-B) Scatter plots showing protein intensity correlation in the nsP3-mScarlet (nsP3-Sca) immunoprecipitations in extracts from cells infected with SINV (A) and SINV_nsP3-mScarlet_ (B). Pearson coefficient and protein intensity distribution are shown.

**Figure S4.**
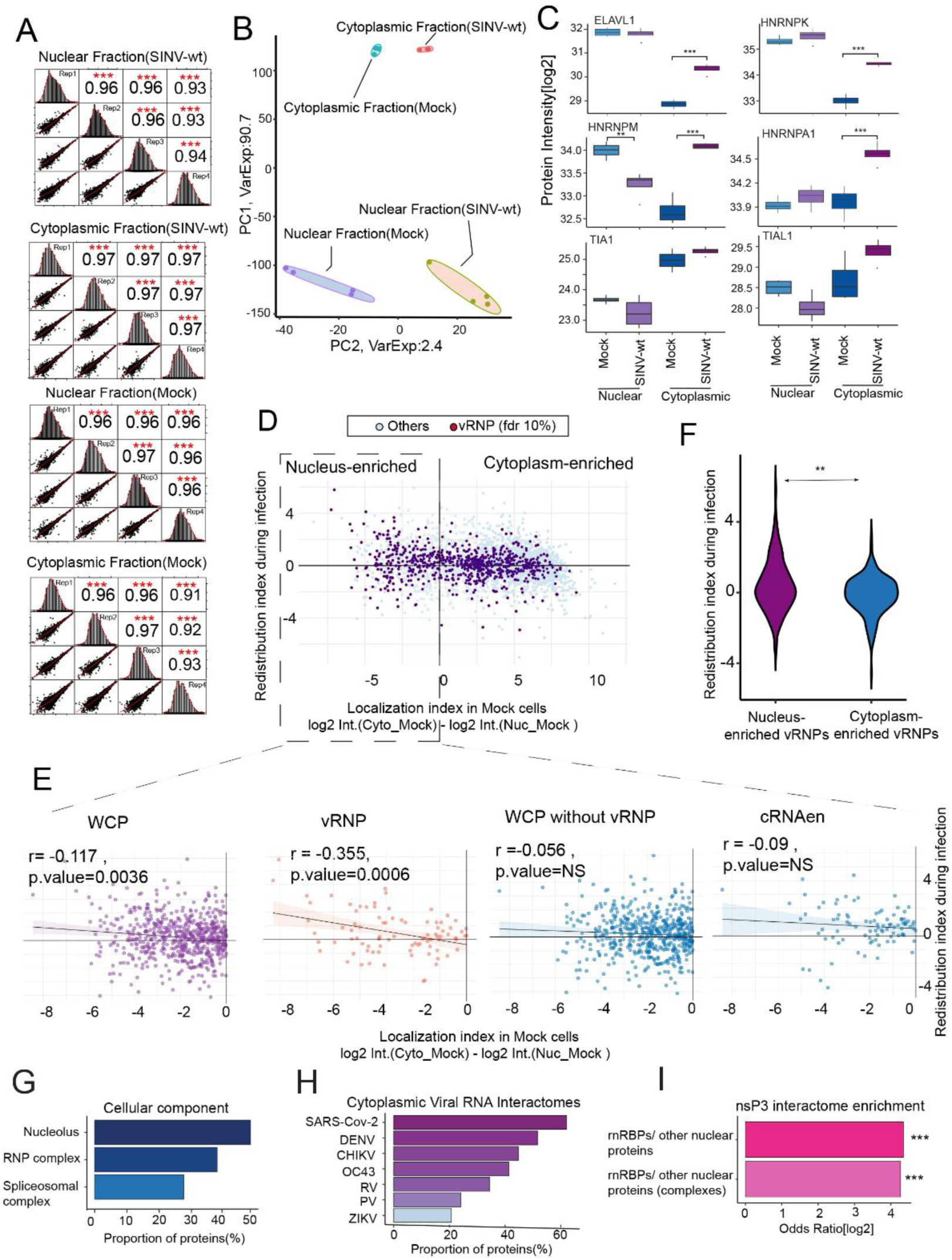
Proteomic analysis of nucleus and cytoplasm in SINV infected cells. A) Scatter plots showing protein intensity correlation in nuclear and cytoplasmic fractions in mock- and SINV-infected cells. Pearson coefficient and protein intensity distribution are shown. B) Principal component analysis (PCA) of the nuclear and cytoplasmic fractions of mock- and SINV-infected cells. C) Intensities of selected proteins in the nucleus and cytoplasm of mock- and SINV-infected cells. These proteins were previously shown to relocate to the cytoplasm in alphavirus infected cells. *, ** and *** indicate FDR < 10, 5 and 1% respectively. D) Scatter plot comparing the localisation index generated in uninfected cells to the redistribution index (localisation index in infected - localisation index mock cells). E) Linear regression analysis of localisation index against redistribution index for following protein subgroups: whole cell proteome (WCP), vRNPs, WCP without vRNPs components, and cRNPen. F) Violin plot showing the distribution of the redistribution index across the proteins within nuclear and cytoplasmic components of vRNPs. ** indicate p-value <0.05 estimated by Welch’s t-test. G) Bar plot showing ‘cellular component’ GO terms enriched in the nuclear components of vRNPs displaying cytoplasmic redistribution in infected cells. H) Proportion of redistributed rnRBPs that interact with SINV RNA (here) and have been reported to interact with the RNAs of other cytoplasmic viruses. Data compiled extracted from (Iselin et al., 2022). I) Bar plot defining the enrichment of rnRBPs or their associated complexes in the nsP3 interactome, curated human complexes obtained from CORUM database (Giurgiu et al., 2019). *** indicate p-value <0.01 estimated by Fisher’s Exact test.

**Figure S5.**
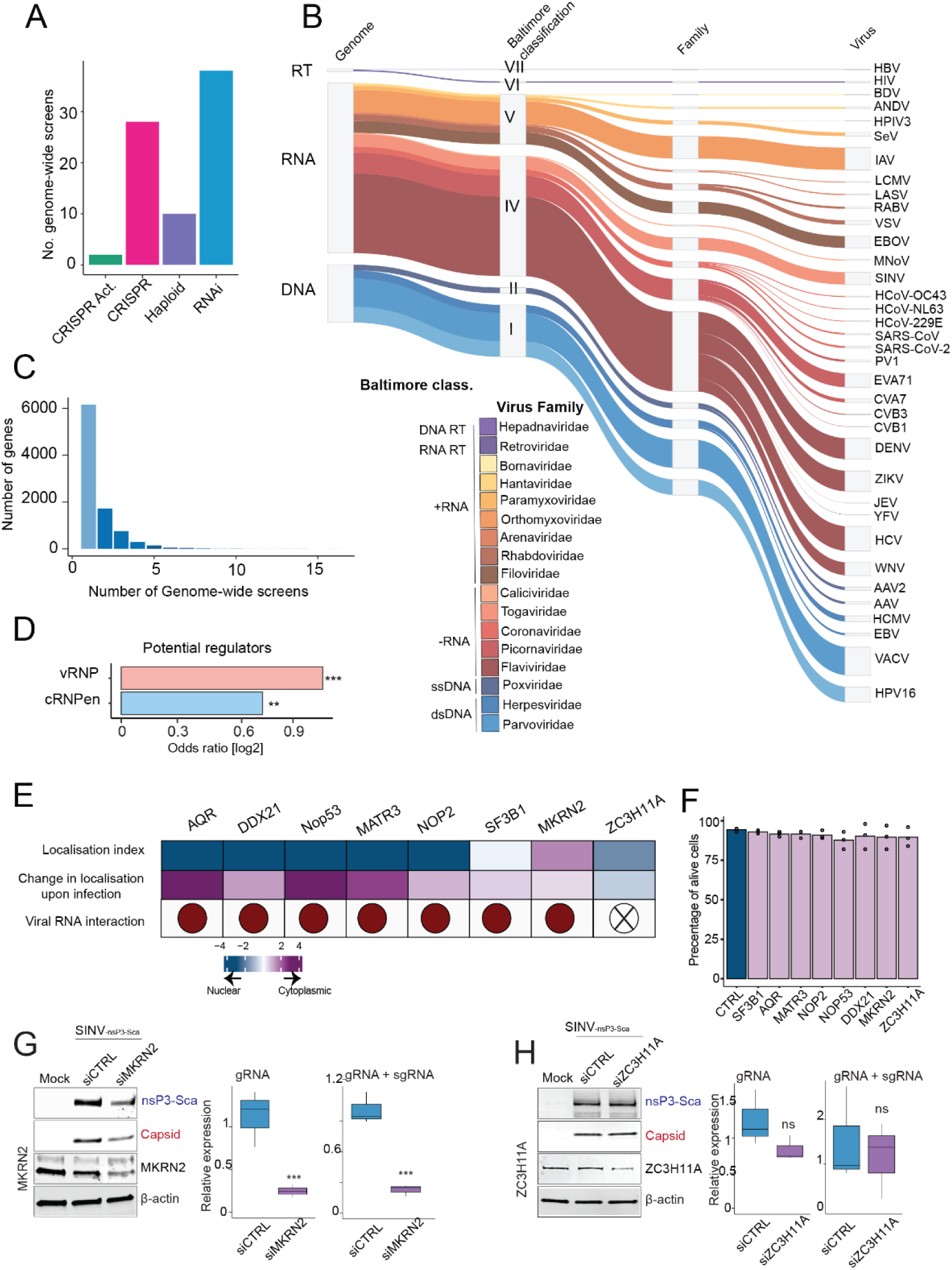
Loss of function of selected rnRBPs affects viral gene expression. A) Bar plot summarising the number of studies for each genome-wide screening strategy. B) Outline of the Baltimore classification of the different viruses covered in the genome-wide screens. Line thickness relates to the total number of reported “hits” (genes). C) Bar plot showing the relationship between genes and number of genome-wide screens reporting them as ‘hits’. D) Bar plot showing the enrichment of potential regulators (proteins reported in three different genome-wide screens) in vRNP or cRNPen datasets. ** and *** indicate p-value <0.05 and <0.01 respectively estimated by Welch’s t-test. E) Properties of the selected candidates for loss of function experiments, including their localisation in uninfected cells, magnitude of change in localisation upon infection (data from Figure 4) and vRNA interaction (X indicate no interaction detected, data from Figure 2). F) Percentage of alive HEK293 cells measured by staining with trypan blue after 24 hours of transfection of the non-targeting siRNA (siCTRL) and specific siRNAs G–H) HEK293 cells treated with non-targeting siRNA (siCTRL) or siRNA against protein of interest (siPOI) for 24 hours, followed by infection with SINV_nsP3-mScarlet_ (MOI = 0.5) for 18 hpi. Left panels,Immunoblotting analysis. Right panel, qRT-PCR analysis with primers against the genomic (gRNA) or genomic and sub-genomic RNA (gRNA + sgRNA), n=3. *, ** and *** indicate student t-test p-value < 0.1, 0.05 and 0.01 respectively.

**Figure S6.**
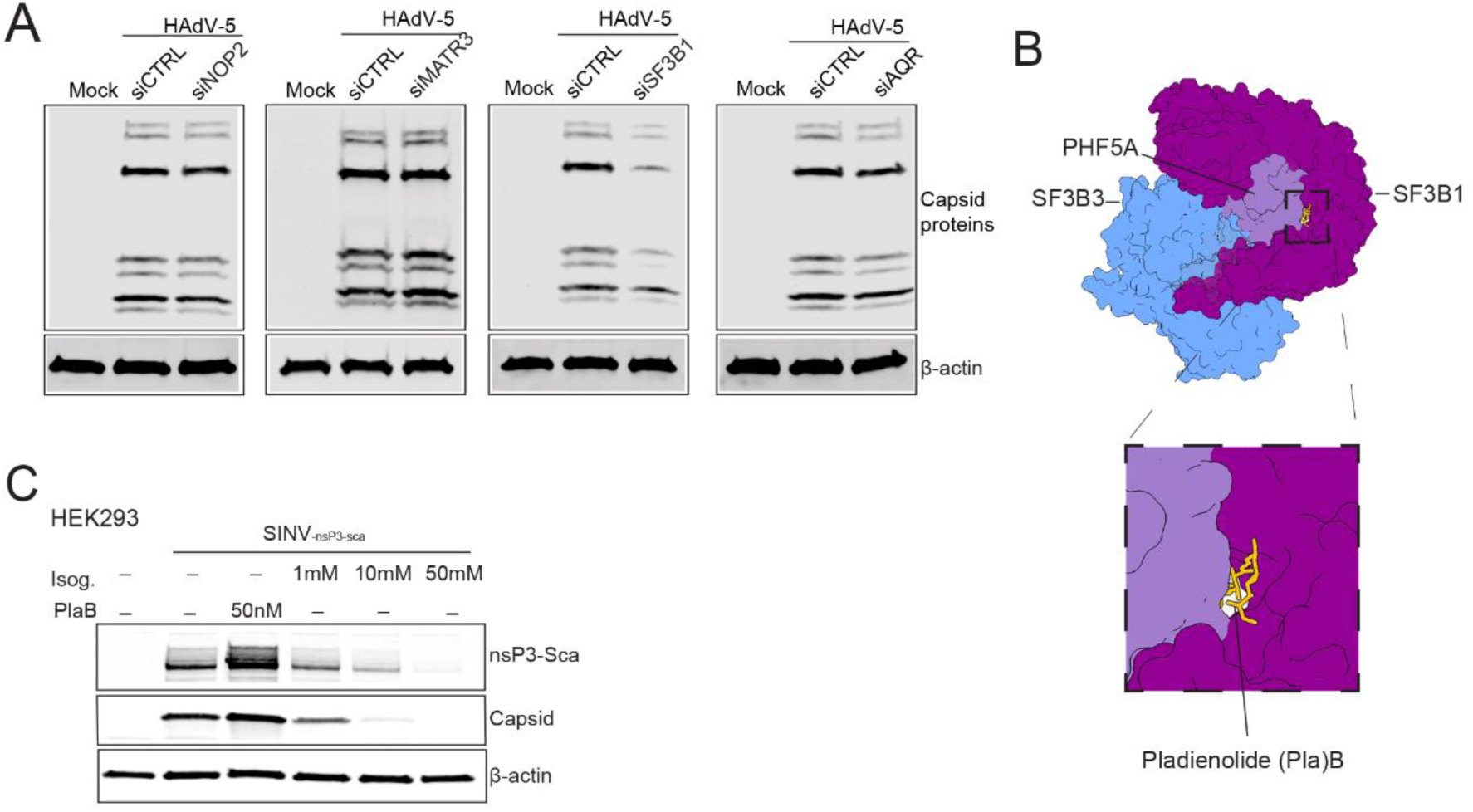
Relevance of vRNP components in virus infection. A) HEK293 cells treated with siRNA against nontargeting (siCTRL) or protein of interest (siPOI) for 48 hours, followed by infection with HAdV-5 (MOI= 1, 24hpi). B) Crystal structure of SF3b core in complex with a splicing modulator PlaB (showing SF3B1, SF3B3, PH5FA and PlaB; PDB ID: 6EN4). C) Immunoblotting analysis of nsP3-Scarlet (nsP3-Sca), capsid and β-actin from mock or SINV_nsP3-mScarlet_ (MOI= 0.1) infected HEK293 cells, pre-treated for 2 hours with DMSO or Pladienolide B (PlaB) or Isoginkgetin (Isog.) at the indicated concentrations.

## REFERENCES

Baltz, A.G., Munschauer, M., Schwanhausser, B., Vasile, A., Murakawa, Y., Schueler, M., Youngs, N., Penfold-Brown, D., Drew, K., Milek, M., et al. (2012). The mRNA-bound proteome and its global occupancy profile on protein-coding transcripts. Mol Cell 46, 674–690.

Barchiesi, A., and Vascotto, C. (2019). Transcription, Processing, and Decay of Mitochondrial RNA in Health and Disease. Int J Mol Sci 20.

Bonizzoni, M., Gasperi, G., Chen, X., and James, A.A. (2013). The invasive mosquito species Aedes albopictus: current knowledge and future perspectives. Trends Parasitol 29, 460–468.

Carrasco, L., Sanz, M.A., and Gonzalez-Almela, E. (2018). The Regulation of Translation in Alphavirus-Infected Cells. Viruses 10.

Castello, A., Fischer, B., Eichelbaum, K., Horos, R., Beckmann, B.M., Strein, C., Davey, N.E., Humphreys, D.T., Preiss, T., Steinmetz, L.M., et al. (2012). Insights into RNA biology from an atlas of mammalian mRNA-binding proteins. Cell 149, 1393–1406.

Castello, A., Fischer, B., Frese, C.K., Horos, R., Alleaume, A.M., Foehr, S., Curk, T., Krijgsveld, J., and Hentze, M.W. (2016). Comprehensive Identification of RNA-Binding Domains in Human Cells. Mol Cell 63, 696–710.

Castello, A., Horos, R., Strein, C., Fischer, B., Eichelbaum, K., Steinmetz, L.M., Krijgsveld, J., and Hentze, M.W. (2013). System-wide identification of RNA-binding proteins by interactome capture. Nat Protoc 8, 491–500.

Castello, A., Sanz, M.A., Molina, S., and Carrasco, L. (2006). Translation of Sindbis virus 26S mRNA does not require intact eukariotic initiation factor 4G. J Mol Biol 355, 942–956.

Cox, J., Neuhauser, N., Michalski, A., Scheltema, R.A., Olsen, J.V., and Mann, M. (2011). Andromeda: a peptide search engine integrated into the MaxQuant environment. J Proteome Res 10, 1794–1805.

Damle, N.P., and Kohn, M. (2019). The human DEPhOsphorylation Database DEPOD: 2019 update. Database (Oxford) 2019.

Dhir, A., Dhir, S., Borowski, L.S., Jimenez, L., Teitell, M., Rotig, A., Crow, Y.J., Rice, G.I., Duffy, D., Tamby, C., et al. (2018). Mitochondrial double-stranded RNA triggers antiviral signalling in humans. Nature 560, 238–242.

Doherty, L.M., Mills, C.E., Boswell, S.A., Liu, X., Hoyt, C.T., Gyori, B., Buhrlage, S.J., and Sorger, P.K. (2022). Integrating multi-omics data reveals function and therapeutic potential of deubiquitinating enzymes. Elife 11.

Effenberger, K.A., Urabe, V.K., and Jurica, M.S. (2017). Modulating splicing with small molecular inhibitors of the spliceosome. Wiley Interdiscip Rev RNA 8.

Flynn, R.A., Belk, J.A., Qi, Y., Yasumoto, Y., Wei, J., Alfajaro, M.M., Shi, Q., Mumbach, M.R., Limaye, A., DeWeirdt, P.C., et al. (2021). Discovery and functional interrogation of SARS-CoV-2 RNA-host protein interactions. Cell 184, 2394–2411 e2316.

Fukuchi, S., Hosoda, K., Homma, K., Gojobori, T., and Nishikawa, K. (2011). Binary classification of protein molecules into intrinsically disordered and ordered segments. BMC Struct Biol 11, 29.

Fukuhara, T., Hosoya, T., Shimizu, S., Sumi, K., Oshiro, T., Yoshinaka, Y., Suzuki, M., Yamamoto, N., Herzenberg, L.A., Herzenberg, L.A., et al. (2006). Utilization of host SR protein kinases and RNA-splicing machinery during viral replication. Proc Natl Acad Sci U S A 103, 11329–11333.

Gao, Y., Goonawardane, N., Ward, J., Tuplin, A., and Harris, M. (2019). Multiple roles of the non-structural protein 3 (nsP3) alphavirus unique domain (AUD) during Chikungunya virus genome replication and transcription. PLoS Pathog 15, e1007239.

Garcia-Moreno, M., Noerenberg, M., Ni, S., Jarvelin, A.I., Gonzalez-Almela, E., Lenz, C.E., Bach-Pages, M., Cox, V., Avolio, R., Davis, T., et al. (2019). System-wide Profiling of RNA-Binding Proteins Uncovers Key Regulators of Virus Infection. Mol Cell 74, 196–211 e111.

Garcia-Moreno, M., Sanz, M.A., Pelletier, J., and Carrasco, L. (2013). Requirements for eIF4A and eIF2 during translation of Sindbis virus subgenomic mRNA in vertebrate and invertebrate host cells. Cell Microbiol 15, 823–840.

Gebhart, N.N., Hardy, R.W., and Sokoloski, K.J. (2020). Comparative analyses of alphaviral RNA:Protein complexes reveals conserved host-pathogen interactions. PLoS One 15, e0238254.

Girard, C., Will, C.L., Peng, J., Makarov, E.M., Kastner, B., Lemm, I., Urlaub, H., Hartmuth, K., and Luhrmann, R. (2012). Post-transcriptional spliceosomes are retained in nuclear speckles until splicing completion. Nat Commun 3, 994.

Giurgiu, M., Reinhard, J., Brauner, B., Dunger-Kaltenbach, I., Fobo, G., Frishman, G., Montrone, C., and Ruepp, A. (2019). CORUM: the comprehensive resource of mammalian protein complexes-2019. Nucleic Acids Res 47, D559–D563.

Gorchakov, R., Frolova, E., and Frolov, I. (2005). Inhibition of transcription and translation in Sindbis virus-infected cells. J Virol 79, 9397–9409.

Gotte, B., Liu, L., and McInerney, G.M. (2018). The Enigmatic Alphavirus Non-Structural Protein 3 (nsP3) Revealing Its Secrets at Last. Viruses 10.

Gutierrez-Chamorro, L., Felip, E., Ezeonwumelu, I.J., Margeli, M., and Ballana, E. (2021). Cyclin-dependent Kinases as Emerging Targets for Developing Novel Antiviral Therapeutics. Trends Microbiol 29, 836–848.

Hobor, F., Dallmann, A., Ball, N.J., Cicchini, C., Battistelli, C., Ogrodowicz, R.W., Christodoulou, E., Martin, S.R., Castello, A., Tripodi, M., et al. (2018). A cryptic RNA-binding domain mediates Syncrip recognition and exosomal partitioning of miRNA targets. Nat Commun 9, 831.

Hughes, C.S., Moggridge, S., Muller, T., Sorensen, P.H., Morin, G.B., and Krijgsveld, J. (2019). Single-pot, solid-phase-enhanced sample preparation for proteomics experiments. Nat Protoc 14, 68–85.

Hull, C.M., and Bevilacqua, P.C. (2016). Discriminating Self and Non-Self by RNA: Roles for RNA Structure, Misfolding, and Modification in Regulating the Innate Immune Sensor PKR. Acc Chem Res 49, 1242–1249.

Iselin, L., Palmalux, N., Kamel, W., Simmonds, P., Mohammed, S., and Castello, A. (2021). Uncovering viral RNA-host cell interactions on a proteome-wide scale. Trends Biochem Sci.

Iselin, L., Palmalux, N., Kamel, W., Simmonds, P., Mohammed, S., and Castello, A. (2022). Uncovering viral RNA-host cell interactions on a proteome-wide scale. Trends Biochem Sci 47, 23–38.

Jarvelin, A.I., Noerenberg, M., Davis, I., and Castello, A. (2016). The new (dis)order in RNA regulation. Cell Commun Signal 14, 9.

Kamel, W., Noerenberg, M., Cerikan, B., Chen, H., Jarvelin, A.I., Kammoun, M., Lee, J.Y., Shuai, N., Garcia-Moreno, M., Andrejeva, A., et al. (2021). Global analysis of protein-RNA interactions in SARS-CoV-2-infected cells reveals key regulators of infection. Mol Cell 81, 2851–2867 e2857.

Kim, B., Arcos, S., Rothamel, K., Jian, J., Rose, K.L., McDonald, W.H., Bian, Y., Reasoner, S., Barrows, N.J., Bradrick, S., et al. (2020). Discovery of Widespread Host Protein Interactions with the Pre-replicated Genome of CHIKV Using VIR-CLASP. Mol Cell 78, 624–640 e627.

Kim, D.Y., Reynaud, J.M., Rasalouskaya, A., Akhrymuk, I., Mobley, J.A., Frolov, I., and Frolova, E.I. (2016). New World and Old World Alphaviruses Have Evolved to Exploit Different Components of Stress Granules, FXR and G3BP Proteins, for Assembly of Viral Replication Complexes. PLoS Pathog 12, e1005810.

Kurkela, S., Manni, T., Vaheri, A., and Vapalahti, O. (2004). Causative agent of Pogosta disease isolated from blood and skin lesions. Emerg Infect Dis 10, 889–894.

Kyriakis, J.M., and Avruch, J. (2012). Mammalian MAPK signal transduction pathways activated by stress and inflammation: a 10-year update. Physiol Rev 92, 689–737.

Laine, M., Luukkainen, R., and Toivanen, A. (2004). Sindbis viruses and other alphaviruses as cause of human arthritic disease. J Intern Med 256, 457–471.

Lanke, K.H., van der Schaar, H.M., Belov, G.A., Feng, Q., Duijsings, D., Jackson, C.L., Ehrenfeld, E., and van Kuppeveld, F.J. (2009). GBF1, a guanine nucleotide exchange factor for Arf, is crucial for coxsackievirus B3 RNA replication. J Virol 83, 11940–11949.

LaPointe, A.T., Gebhart, N.N., Meller, M.E., Hardy, R.W., and Sokoloski, K.J. (2018). The Identification and Characterization of Sindbis Virus RNA:Host Protein Interactions. J Virol.

Lazar, C., Gatto, L., Ferro, M., Bruley, C., and Burger, T. (2016). Accounting for the Multiple Natures of Missing Values in Label-Free Quantitative Proteomics Data Sets to Compare Imputation Strategies. J Proteome Res 15, 1116–1125.

Lee, A.S., Kranzusch, P.J., Doudna, J.A., and Cate, J.H. (2016). eIF3d is an mRNA cap-binding protein that is required for specialized translation initiation. Nature 536, 96–99.

Lee, J.Y., Wing, P.A.C., Gala, D.S., Noerenberg, M., Jarvelin, A.I., Titlow, J., Zhuang, X., Palmalux, N., Iselin, L., Thompson, M.K., et al. (2022). Absolute quantitation of individual SARS-CoV-2 RNA molecules provides a new paradigm for infection dynamics and variant differences. Elife 11.

Lee, S., Kim, J.Y., Kim, Y.J., Seok, K.O., Kim, J.H., Chang, Y.J., Kang, H.Y., and Park, J.H. (2012). Nucleolar protein GLTSCR2 stabilizes p53 in response to ribosomal stresses. Cell Death Differ 19, 1613–1622.

Lee, S., Lee, Y.S., Choi, Y., Son, A., Park, Y., Lee, K.M., Kim, J., Kim, J.S., and Kim, V.N. (2021). The SARS-CoV-2 RNA interactome. Mol Cell 81, 2838–2850 e2836.

Lim, E.X.Y., Lee, W.S., Madzokere, E.T., and Herrero, L.J. (2018). Mosquitoes as Suitable Vectors for Alphaviruses. Viruses 10.

Lin, X., Zhou, L., Zhong, J., Zhong, L., Zhang, R., Kang, T., and Wu, Y. (2022). RNA-binding protein RBM28 can translocate from the nucleolus to the nucleoplasm to inhibit the transcriptional activity of p53. J Biol Chem 298, 101524.

Lloyd, R.E. (2015). Nuclear proteins hijacked by mammalian cytoplasmic plus strand RNA viruses. Virology 479-480, 457–474.

Manning, G., Whyte, D.B., Martinez, R., Hunter, T., and Sudarsanam, S. (2002). The protein kinase complement of the human genome. Science 298, 1912–1934.

Marks, M., and Marks, J.L. (2016). Viral arthritis. Clin Med (Lond) 16, 129–134.

O’Brien, K., Matlin, A.J., Lowell, A.M., and Moore, M.J. (2008). The biflavonoid isoginkgetin is a general inhibitor of Pre-mRNA splicing. J Biol Chem 283, 33147–33154.

Park, E., and Griffin, D.E. (2009). Interaction of Sindbis virus non-structural protein 3 with poly(ADP-ribose) polymerase 1 in neuronal cells. J Gen Virol 90, 2073–2080.

Patricio, D.O., Dias, G.B.M., Granella, L.W., Trigg, B., Teague, H.C., Bittencourt, D., Bafica, A., Zanotto-Filho, A., Ferguson, B., and Mansur, D.S. (2022). DNA-PKcs restricts Zika virus spreading and is required for effective antiviral response. Front Immunol 13, 1042463.

Perez-Perri, J.I., Noerenberg, M., Kamel, W., Lenz, C.E., Mohammed, S., Hentze, M.W., and Castello, A. (2021). Global analysis of RNA-binding protein dynamics by comparative and enhanced RNA interactome capture. Nat Protoc 16, 27–60.

Peters, N.E., Ferguson, B.J., Mazzon, M., Fahy, A.S., Krysztofinska, E., Arribas-Bosacoma, R., Pearl, L.H., Ren, H., and Smith, G.L. (2013). A mechanism for the inhibition of DNA-PK-mediated DNA sensing by a virus. PLoS Pathog 9, e1003649.

Piovesan, D., Necci, M., Escobedo, N., Monzon, A.M., Hatos, A., Micetic, I., Quaglia, F., Paladin, L., Ramasamy, P., Dosztanyi, Z., et al. (2021). MobiDB: intrinsically disordered proteins in 2021. Nucleic Acids Res 49, D361–D367.

Sanz, M.A., Garcia-Moreno, M., and Carrasco, L. (2015). Inhibition of host protein synthesis by Sindbis virus: correlation with viral RNA replication and release of nuclear proteins to the cytoplasm. Cell Microbiol 17, 520–541.

Schmidt, N., Lareau, C.A., Keshishian, H., Ganskih, S., Schneider, C., Hennig, T., Melanson, R., Werner, S., Wei, Y., Zimmer, M., et al. (2021). The SARS-CoV-2 RNA-protein interactome in infected human cells. Nat Microbiol 6, 339–353.

Schwarzl, T., Group., H., and Grouo., H. (2020). RBPbase v0.2.1 alpha, EMBL, ed.

Silva, L.A., and Dermody, T.S. (2017). Chikungunya virus: epidemiology, replication, disease mechanisms, and prospective intervention strategies. J Clin Invest 127, 737–749.

Sokoloski, K.J., Nease, L.M., May, N.A., Gebhart, N.N., Jones, C.E., Morrison, T.E., and Hardy, R.W. (2017). Identification of Interactions between Sindbis Virus Capsid Protein and Cytoplasmic vRNA as Novel Virulence Determinants. PLoS Pathog 13, e1006473.

Strauss, J.H., and Strauss, E.G. (1994). The alphaviruses: gene expression, replication, and evolution. Microbiol Rev 58, 491–562.

Sun, C. (2020). The SF3b complex: splicing and beyond. Cell Mol Life Sci 77, 3583–3595.

Tan, Y.B., Chmielewski, D., Law, M.C.Y., Zhang, K., He, Y., Chen, M., Jin, J., and Luo, D. (2022). Molecular architecture of the Chikungunya virus replication complex. Sci Adv 8, eadd2536.

Taylor, S.S., Haste, N.M., and Ghosh, G. (2005). PKR and eIF2alpha: integration of kinase dimerization, activation, and substrate docking. Cell 122, 823–825.

Temperley, R.J., Wydro, M., Lightowlers, R.N., and Chrzanowska-Lightowlers, Z.M. (2010). Human mitochondrial mRNAs--like members of all families, similar but different. Biochim Biophys Acta 1797, 1081–1085.

Thompson, L., Depledge, D.P., Burgess, H.M., and Mohr, I. (2022). An eIF3d-dependent switch regulates HCMV replication by remodeling the infected cell translation landscape to mimic chronic ER stress. Cell Rep 39, 110767.

Van Nostrand, E.L., Pratt, G.A., Shishkin, A.A., Gelboin-Burkhart, C., Fang, M.Y., Sundararaman, B., Blue, S.M., Nguyen, T.B., Surka, C., Elkins, K., et al. (2016). Robust transcriptome-wide discovery of RNA-binding protein binding sites with enhanced CLIP (eCLIP). Nat Methods 13, 508–514.

Ventoso, I., Sanz, M.A., Molina, S., Berlanga, J.J., Carrasco, L., and Esteban, M. (2006). Translational resistance of late alphavirus mRNA to eIF2alpha phosphorylation: a strategy to overcome the antiviral effect of protein kinase PKR. Genes Dev 20, 87–100.

Wang, C., Chua, K., Seghezzi, W., Lees, E., Gozani, O., and Reed, R. (1998). Phosphorylation of spliceosomal protein SAP 155 coupled with splicing catalysis. Genes Dev 12, 1409–1414.

Westergren Jakobsson, A., Segerman, B., Wallerman, O., Lind, S.B., Zhao, H., Rubin, C.J., Pettersson, U., and Akusjarvi, G. (2021). The Human Adenovirus Type 2 Transcriptome: An Amazing Complexity of Alternatively Spliced mRNAs. J Virol 95.

Yang, K., Wang, M., Zhao, Y., Sun, X., Yang, Y., Li, X., Zhou, A., Chu, H., Zhou, H., Xu, J., et al. (2016). A redox mechanism underlying nucleolar stress sensing by nucleophosmin. Nat Commun 7, 13599.

Younis, S., Kamel, W., Falkeborn, T., Wang, H., Yu, D., Daniels, R., Essand, M., Hinkula, J., Akusjarvi, G., and Andersson, L. (2018). Multiple nuclear-replicating viruses require the stress-induced protein ZC3H11A for efficient growth. Proc Natl Acad Sci U S A 115, E3808–E3816.

